# Size photometry and fluorescence imaging of immobilized immersed extracellular vesicles

**DOI:** 10.1101/2023.12.08.570449

**Authors:** Andreas Wallucks, Philippe DeCorwin-Martin, Molly L. Shen, Andy Ng, David Juncker

## Abstract

Immunofluorescence analysis of individual extracellular vesicles (EVs) in common fluorescence microscopes is gaining popularity due to its accessibility and high fluorescence sensitivity, however, EV number and size are only measurable using fluorescent stains requiring extensive sample manipulations. Here we introduce highly sensitive label-free EV size photometry (SP) based on interferometric scattering (iSCAT) imaging of immersed EVs immobilized on a glass coverslip. We implement SP on a common inverted epifluorescence microscope with LED illumination and a simple 50:50 beamsplitter, permitting seamless integration of SP with fluorescence imaging (SPFI). We present a high-throughput SPFI workflow recording >10,000 EVs in 7 min over multiple fields of view, pre- and post-incubation imaging to suppress background, along with automated image alignment, aberration correction, spot detection, and EV sizing. We identify the upper sizing limit of SP with 440 nm illumination and extend EVs sizing from ∼37 nm in diameter to >200 nm with dual 440 nm and 740 nm illumination. We benchmark SP to flow cytometry using calibrated silica nanoparticles and demonstrate superior, label-free sensitivity. We showcase SPFI’s potential for EV analysis by experimentally distinguishing surface and volumetric EV dyes, observing the deformation of EVs adsorbed to a surface, and by uncovering distinct subpopulations in <100 nm-in-diameter EVs with fluorescently tagged membrane proteins.

## INTRODUCTION

Cells secrete membrane-bound structures called extracellular vesicles (EVs) that are fundamental to tissue homeostasis and cell signaling^1^, and have garnered great interest as biomarkers and for therapeutics^2^. Their sizes range from a few tens of nanometers to more than a micrometer, and they carry a large variety of bioactive molecules such as proteins, lipids, nucleic acids, and metabolites. Major efforts have been devoted to the development of single vesicle measurement techniques to characterize the heterogeneity of EVs and the features that govern cellular activation pathways, EV cargo sorting^3^ and how they change in health and disease^4^. Currently, there are no generally accepted classifiers to distinguish EV subpopulations such as exosomes or ectosomes due to various technical challenges. Single protein expression levels suggest EV subpopulations, but due to considerable overlap are insufficient for classification^5^. Multi-dimensional characterization will be required and could involve multiplex proteomic analysis or incorporation of orthogonal parameters such as EV size. It is furthermore understood that EV diameters in various samples follow inverse power-law distributions where small EVs with diameter *d* < 100 nm are by far the most abundant^6,7^. Consequently, the limit of detection (LOD) of any single EV technology profoundly influences the results. For instance, proteomic analysis significantly varies between measurements with LODs of 100 nm compared to 150 nm^8^, suggesting that techniques with LODs substantially below 100 nm are required for reproducible and unbiased sample characterization.

Several techniques show potential to meet these requirements and advance EV subclassification. For example, electron microscopy can be combined with immunogold labelling to both size and characterize EVs down to *d* < 30 nm but has exceedingly low throughput and is thus impractical. Optical methods that can combine EV immunofluorescence analysis with EV sizing based on light scattering are preferred owing to their higher throughput. Indeed, flow cytometry (FCM) emerged as the current gold standard for single EV analysis owing to its comparative ease of use, low operating cost, and simultaneous measurement of both EV size (by side scattering) and protein expression (by immunofluorescence)^8^. Many common flow cytometers used to detect EVs have an LOD ranging between *d* ≈ 100 – 150 nm, while specialized nanoflow cytometers can detect EVs as small as *d* ≈ 40 nm, but at the expense of fluorescence multiplexing^9^. The LOD of FCM is commensurate to nanoparticle tracking analysis (NTA) with a label-free LOD of *d* ≈ 80 nm which is commonly used to assess EV size distributions. Lowering the detection threshold remains a challenge as nanoparticle scattering scales with *d*^!^, and lowering the LOD just three-fold requires almost a thousand-fold gain in sensitivity. Recently, fluorescence imaging of EVs captured on a surface by widefield or TIRF microscopy^10–12^ has gained traction as an alternative to FCM due to its accessibility, higher fluorescence sensitivity^13^, and the possibility to manipulate EVs following their immobilization^14^. The absence of a scatter signal for sizing is circumvented using fluorescent labels (e.g. dyes), but at the cost of further sample manipulation and susceptibility to false positive detection events, e.g. due to label aggerates.

Alternatively, super-resolution imaging can be used to size EVs down to *d* ≈ 30 nm but suffers from low throughput and requires very high expression of target proteins (at least 20) in a single EV^6^. Label-free detection of surface bound particles by dark-field imaging is well established^15^, however, it requires specialized microscopes with bright light sources and strong stray light suppression, and the scattering signal also scales unfavorably ∝ *d*^6^, and has not been useful for EV analysis.

Interference imaging methods such as interference scattering microscopy (iSCAT)^16^, single particle Interference Reflection Image Sensing (SP-IRIS)^17^, and several others^18–21^ offer relief as they optically amplify signals with minimally added noise^22^, are robust against stray light, and benefit from a more favorable contrast scaling ∝ *d*^3^. iSCAT and SP-IRIS require the least complex optical setups as both generate the interferometric reference field by reflection of the illumination light from the imaging substrate in common-path geometry. SP-IRIS achieves this using custom-made silicon chips coated with a thin dielectric reflective layer. It employs imaging from the top, mostly though air, since immersing samples adds complications such as additional reflections from the top surface or minute changes in liquid depth that affect focusing. iSCAT on the other hand uses common glass coverslips without any modification. Imaging is conducted not through the medium, but through the coverslip, and immersed samples can readily be analyzed. iSCAT offers exceptional sensitivity including label-free single molecule detection and mass photometry^23^, it can be combined with fluorescence^24^ and has been used to detect EVs^25,26^. However, most iSCAT setups use specialized hardware such as high-powered lasers and high-speed cameras which are not tailored for wide-field EV fluorescence analysis over large fields of view, so far limiting the adoption of iSCAT for EV research.

Here we introduce EV size photometry (SP) using iSCAT on a standard widefield fluorescence microscope with non-imaging based autofocus by adding only a simple 50:50 beamsplitter and by reusing LED light source and sCMOS camera intended for fluorescence microscopy. We demonstrate sizing of various immobilized nanoparticles by pre- and post-incubation imaging of a capture surface and benchmark it against flow cytometry using calibrated silica particles.

Recognizing an upper sizing limit for SP, we introduce combined SP using two illumination wavelengths and extend the dynamic range of SP for polydisperse samples such as liposomes and EVs from *d* ≈ 37 to > 200 nm. Our analysis is facilitated by an open-source automated high throughput image analysis pipeline that includes image alignment, background suppression, and aberration correction steps to register and analyse captured particles over multiple fields of view (FOVs). We demonstrate the natural integration of SP with highly sensitive EV fluorescence imaging (SPFI) and validate the sizing accuracy on EV samples stained with surface and intraluminal dyes as function of EV diameter by correlating SP contrasts with fluorescence intensities. We furthermore observe SP contrast behavior consistent with EV deformation on the surface. Finally, we analyze EVs expressing a fluorescent fusion protein and identify two distinct clusters of EVs with *d* < 100 nm, which are revealed by correlating EV sizes with the expression of the protein tag and would be missed by techniques with LODs of *d* > 100 nm.

## RESULTS

SPFI allows to combine sizing of immersed EVs inside a microfluidic flow cell with further characterization using fluorescent reporters or affinity binders. Particle sizing is enabled through label free iSCAT imaging on common glass coverslips which can be implemented in any widefield epifluorescence microscope by the addition of a 50:50 beamsplitter that is switched with the dichroic filter cube, shown in Fig. 1**a**. We use the terminology ‘size photometry’ because nanoparticles are conventionally characterized by size, however, we note that iSCAT contrasts are principally dependent on both EV size and refractive index and only few implementations truly decouple them^25–27^. Throughout this manuscript, we validate the SP *sizing* accuracy for multiple types of large nanoparticles with increasing complexity (various solid nanoparticles, liposomes, and EVs). A detailed description of the iSCAT imaging principle is given in Supplementary Section S1, describing the mathematical underpinning of its increased sensitivity for small particles compared to darkfield imaging. SP records pairs of pre-incubation (*F*_pre_) and post-incubation (*F*_post_) images over multiple FOVs to maximize imaging throughput while being selective to particles that are immobilized on the coverslip during incubation, Fig. 1**b**. Each *F*_pre_ and *F*_post_contain only static features and can be averaged indefinitely, Fig. 1**d**, relaxing the need for high speed cameras which are otherwise common for dynamic iSCAT imaging^28,29^.

**Figure 1.**
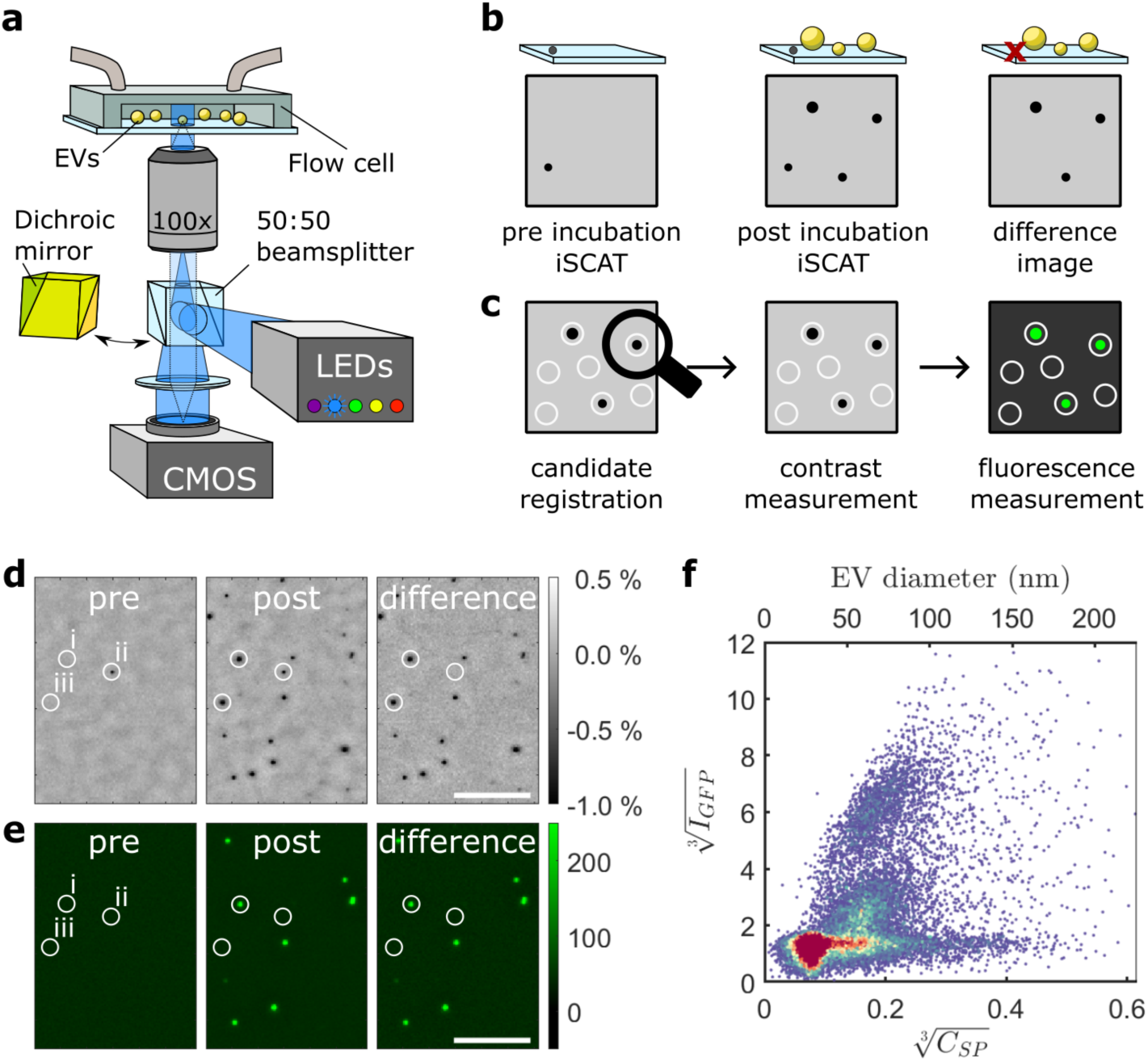
Overview of combined EV size photometry (SP) and fluorescence imaging (SPFI). **a** Schematic of the microscopy setup including a microfluidic flow cell with EVs immobilized on a coverslip in an epifluorescence microscope with LED illumination, a high sensitivity sCMOS camera, and an off-the-shelf 50:50 beamsplitter for iSCAT imaging. A motorized filter turret automatically switches between the dichroic mirror for fluorescence imaging, and the 50:50 beam splitter for iSCAT. **b** Workflow to suppress background inhomogeneities and surface bound crud. The background is registered in a pre-incubation iSCAT image, then EVs are immobilized, unbound EVs are washed, and a post incubation iSCAT image is taken. The difference of the two images efficiently removes small surface inhomogeneities and allows to measure accurate contrast values of small particles within an undulating background. **c** Workflow to generate SPFI scatterplots. Typically, the iSCAT image is used to register candidate spots and contrasts or fluorescence intensities are subsequently measured on all channels. **d** differential iSCAT of pre- and post-incubation image which is used for EV localization. pre: image before particle incubation; post: image after particle incubation; SP: pre-image subtracted from the post-image. Three particles are highlighted: i is a particle that is already present before incubation and is excluded from the rest of the analysis; ii is a particle that is detected in fluorescence, but not in iSCAT, and iii is a particle that is detected both in iSCAT and in fluorescence. **E** Fluorescence pre- and post-incubation images that were taken concomitantly to the iSCAT in **d.** The generation of a difference image in fluorescence is optional, and mostly inconsequential for substrates with low autofluorescence. **f** SPFI scatterplot of the processed data shown in **d** and **e**, taken on EVs from A431 cells that were stably transfected to express CD63 fused with green fluorescence protein.

We introduce an SPFI analysis pipeline (https://github.com/junckerlab/SPFI) to handle data pre-processing and image analysis starting with a step to correct chromatic aberrations between different imaging channels to maximize FOV sizes while ensuring correct spot matching between the channels (Supplementary Section S2). Our calibration-free algorithm can map and correct distortions between any two SPFI channels directly on the respective post-incubation images. Next, the algorithm digitally aligns *F*_pre_and *F*_post_images (Supplementary Section S3) and detects large surface inhomogeneities on *F*_pre_to be excluded from subsequent analysis. Our algorithm then localizes candidate spots in the image data as local contrast minima in *F*_diff_ = *F*_post_ − *F*_pre_, thereby identifying captured EVs as well as sampling the image background to generate an experimental noise floor and inform thresholding (Fig. 1**c**, Supplementary Section S4). SP measurements are taken either in a single illumination wavelength or under dual illumination with a short and a long wavelength to maximise sizing range (described below) and candidate contrasts are measured as center pixel values.

For SPFI, fluorescence images (Fig. 1**e**) are superposed with the iSCAT images and fluorescence intensities *I* for all candidate spots are measured (c.f. Fig. 1**c**, right panel). Taking differential fluorescence images is possible but often inconsequential due to overall low autofluorescence background in empty flow cells. The correlation between SP measurements and fluorescence signal intensities are reported as scatter plots of *I* and *C* (Fig. 1**f**) which are reminiscent of FCM data and led us to adopt data analysis workflows and thresholding protocols common to FCM. EVs are identified in them as candidate spots that exceed the SP background noise found in buffer control experiments, thus establishing a count of the total EV number and removing the ambiguity of thresholding on EV fluorescence alone^12^.

Our analysis pipeline includes an iSCAT flatfield pre-processing step to correct setup-specific illumination inhomogeneities in the raw iSCAT images. The illumination background is identical for each FOV and can vary up to ∼20 % within each image depending on the region of interest (ROI), see Fig. 2**a**, 2**b**. Flatfield processing is necessary because pre- and post-incubation images are often misaligned up to ∼15 % of the FOV size (shifts are caused by manual pipetting and imperfect stage movements) and subtraction of the raw images can lead to residual background that overwhelms EV contrasts <1%. Flatfield images are collected as series of images from random positions over the sample which are averaged into a single flatfield image *FF*. The pre- and post-incubation images are then background corrected by division like *F*_pre_ → *F*_pre_/*FF* and *F*_post_ → *F*_post_/*FF*.

**Figure 2.**
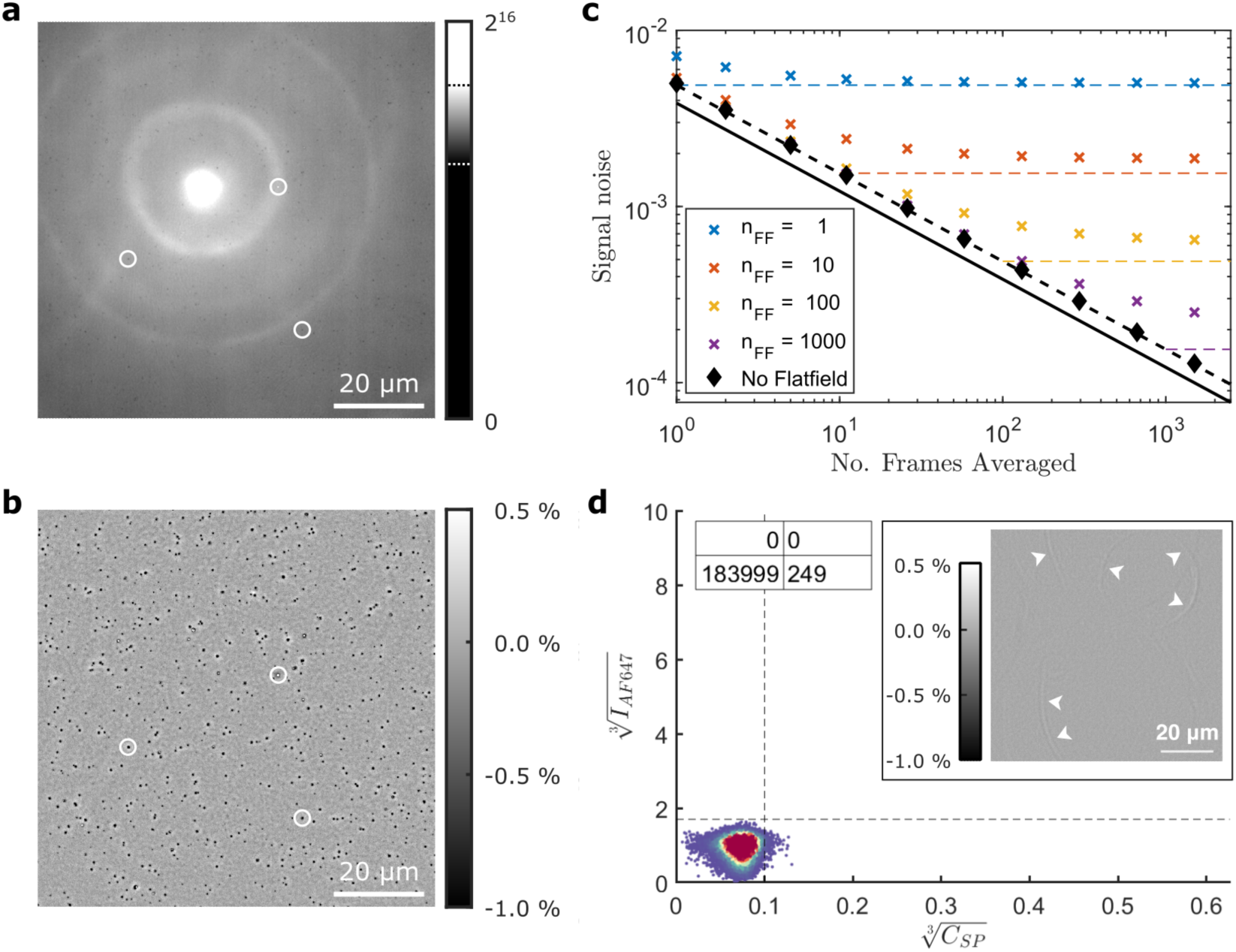
SP noise characterization. **a** Image flatfield processing demonstrated on a raw post-incubation image and **b** flatfield corrected image, which is necessary to remove illumination inhomogeneities but adds image noise. Shown is a 1200×1200 pixel ROI of a raw 16-bit iSCAT image which has close to 20% illumination inhomogeneity. The color bar for the raw image is scaled to the bit depth of the image and dashed lines indicate limits used for plotting. The processed image is shown on a contrast greyscale of 1.5% around the image median. Dark spots are EVs from HT29 cells and circles indicate matching spots as a guide to the eye. **c** iSCAT image noise for a series of images collected at 40 FPS as function of number of images averaged. Black solid line represents an estimate of the photon shot noise and constitutes the lower theoretical noise limit whereas the black dashed line also includes the estimated camera readout noise. The black diamonds are measured SD of the image noise in the averaged timeseries, and crosses are for measured image noise after normalizing the timeseries with flatfield (FF) images consisting of *n*_FF_ frames. The latter is compared to the respective estimated noise floor (horizontal dashed lines) and used to set imaging parameters for the iSCAT endpoint measurements. **d** SPFI plot generated in a buffer control experiment over 20 FOVs using iSCAT at 440 nm wavelength and a red fluorescence channel with vertical gate set three standard deviations above the noise. The plot shows zero localizations in the upper right quadrant and 249 localizations in the bottom right quadrant, many of which stem from setup-specific image artifacts shown in the inset with a differential iSCAT image as white arrowheads.

### iSCAT imaging parameters and image noise

The iSCAT images in this manuscript were taken by averaging 200 frames for both pre- and post-incubation images and normalizing them using flatfield images consisting of 960 individual frames (120 arbitrary positions on a slide to eliminate local interference by dust with 8× average each). We found those imaging parameters by considering the trade-off in imaging speed and detection sensitivity due to image noise as described below. The time it took to capture these images depended on the chosen FOV size since this limited camera acquisition speed and saving time. For example, in 1200×1200 pixel (88 × 88 µm^2^) FOVs with 200 frames could be taken in ∼12 s, split roughly equally between actual camera exposure and software overhead from computing the average and storing the data on the hard disk drive. Flatfield images needed to be taken only once per substrate regardless of the number of imaged FOVs and took about 180 s each. Imaging ten different FOVs in one iSCAT channel with 560 EVs each thus took about 5 min, or 20 FOVs with ∼11,000 EVs total about 7 min (Supplementary Section S5).

The choice of imaging parameters presents a trade-off between imaging time and final image noise, and thus detection sensitivity. The image noise in iSCAT is commonly minimized using cameras with high dynamic range and well depth, exposing pixels close to saturation, as well as averaging of multiple individual frames. Fig. 2**c** shows an image noise characterization for a Prime 95B (Photometrics) with a well depth of 80,000 *e*^-^and readout noise of 1.6 *e*^-^ (see methods). The solid black line indicates the theoretical noise floor consisting purely of photon shot noise (square root of the number of detected photons) whereas the dashed black line includes technical camera readout noise according to manufacturer specifications. This prediction was validated by taking pairs of iSCAT images (each averaged *n* times), subtracting the two to remove fixed-pattern noise, and calculating the standard deviation *σ*_n_ (black diamonds). The noise in SP data includes a second contribution from flatfield processing which adds noise depending on the number of averages *n*_//_ of the flatfield image (crosses, color coded). We found that the measured noise saturated close to the expected noise floor (dashed lines) for small *n*_00_ but exceeded the prediction for larger *n*_00_ slightly. For the imaging parameters used throughout this manuscript, we predicted an approximate image standard deviation *σ* = 5 × 10^-1^ where the flatfield contributed less than a third. The noise floor for SP contrasts *C*_23_exceeded this value (Fig. 2**d** and below, note third root scaling) because our candidate spot localization registers contrast minima and thus samples at the extreme of the noise distribution. This additional noise overhead stemmed from small focus offsets between the averaged flatfield images and the actual FOV due to slow focus drifts which produced artifacts shown in the inset (see also Supplementary Section S6). We calculate our SP LOD from Fig. 2**d** as the mean plus three times the standard deviation (SD) of the noise, resulting in 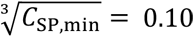. A more stringent definition, for example five SD above the background resulting in 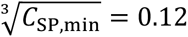, can be chosen for application requiring higher detection specificity.

### Size photometry of silica nanoparticles

We characterized the sensitivity of SP using silica nanoparticles of known sizes and compared it to nanoparticle FCM using a CytoFLEX-S instrument (Beckman Coulter Life Sciences). We measured violet side scattering (VSSC using a 405 nm laser) and side scattering (SSC using a 488 nm laser) of aminated silica particles with 50 nm, 100 nm and 200 nm diameters in Fig. 3**a**. We found that the instrument was able to resolve 100 nm and 200 nm particles which is in line with an ∼84 nm detection limit obtained by the manufacturer on an optimized instrument^30^. Notably, even for the 200 nm particles the signal was within the instrument noise level and could only be distinguished at high sample concentrations and high detection rates.

**Figure 3.**
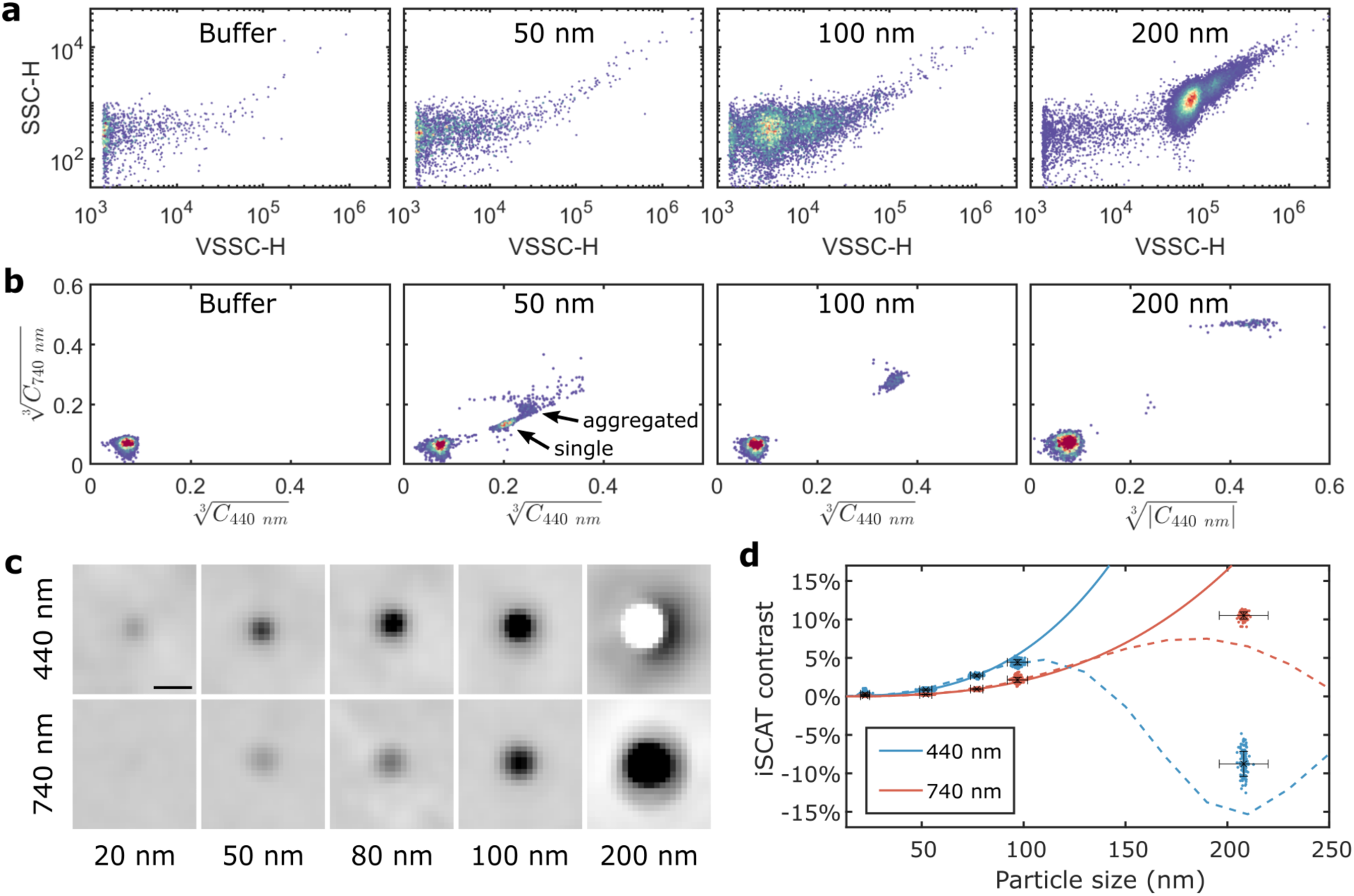
Calibration, validation, and benchmarking of SP. **a** Nanoparticle flow cytometry (FCM) and **b** Size photometry (SP) scatterplots of a buffer control and silica nanoparticles with 50 nm, 100 nm, and 200 nm diameters. For FCM, the samples were diluted to matching concentrations based on the manufacturer specification, measured for 60 s each, and 5 % of total events are shown by their violet side scattering signal (VSSC-H at 405 nm) and side scattering signal (SSC-H at 488 nm). SP plots show iSCAT contrasts to the third root for 440 nm and 740 nm illumination wavelength, respectively. The 50 nm particle signal in SP is clearly separated from the noise blob whereas for FCM no cluster is visible. **c** Representative SP images of particles of different sizes. All color scales are set from -1.0 % to +0.5 %, scalebar is 500 nm. Interference contrast inversion for 200 nm nanoparticles is observed for 440 nm illumination. **d** Theoretical (solid lines: fit to analytical model; dashed lines: calculated by finite element modeling) and experimental (dots) SP contrasts as function of particle size. Horizontal and vertical error bars are SD of the particle sizes specified by the vendor and SD of particle contrast, respectively.

We next took SP measurements under 440 nm and 740 nm iSCAT illumination wavelengths after adsorbing the particles onto plasma-activated glass coverslips. We show SP scatterplots for 50 nm, 100 nm, and 200 nm particles in Fig. 3**b** and representative post-incubation images for these particles as well as for 20 nm and 80 nm ones in Fig. 3**c**. We found that we could detect particles in all samples (Fig. 3**c**) and that the 50 nm particles were the first to cluster separately from the background noise floor (Fig. 3**b**). The scatterplot for the 50 nm particles revealed many particles outside the main cluster which were identified in the raw images as large, non-circular iSCAT spots. These are consistent with aggregates of two or more particles which thus likely dominated the corresponding FCM plot. Similar aggregates were also visible for 100 nm and 200 nm particles but produced excessively bright iSCAT contrasts and were discarded by the localization algorithm. For the 200 nm particles (rightmost panel in Fig. 3**c**), we observed a contrast inversion under 440 nm illumination which requires special attention for SP sizing. The standard mathematical model of interferometric imaging (Supplementary Section 1) predicts that contrasts *C* initially scale with particle diameters *d* to the third power^25^. This *C* ∝ *d*^3^ scaling is an approximation that breaks for large particles, requiring experimental validation to define its range of applicability and establish the maximum sizing range achievable with SP. For the silica standards, we combined measurements *C*_7_ of 20 nm, 50 nm, 80 nm, 100 nm, and 200 nm particles for both wavelengths *λ* in Fig. 3**d**. We fitted the first three samples to *d*^3^ curves (solid lines, color-coded for wavelength), demonstrating that the *d*^3^ scaling extends to 80 nm particles under 400 nm illumination and to 100 nm (but not 200 nm) particles under 740 nm illumination. We continued to investigate the breakdown of the *d*^3^ scaling and contrast inversion of *C*_118 65_ using finite element (FEM) simulations in the COMSOL optics module (Supplementary Section S7). The simulation data, plotted as dashed lines in Fig. 3**d**, reproduced the contrast inversion observed for 200 nm particles with 440 nm illumination and the absence of contrast inversion for 200 nm particles with 740 nm illumination. The simulations furthermore indicate that the *d*^3^ scaling expands to ∼ 90 nm and ∼150 nm particles under 440 nm and 740 nm illumination, respectively. While these simulations improve on the analytical *d*^3^ model, they still miss important effects such as the Gouy phase shift^31^ from microscope focus (Supplementary Section S7) and phase factors stemming from optical aberration in the microscope setup^32^. Future work on analytical treatments of these effects or using machine learning approaches will be instrumental to extend the SP sizing range by using information in the rings around large particles to alleviate ambiguity of contrasts around the inversion point.

We next calibrated SP contrasts for sizing of various nanoparticles including EVs using 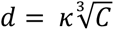, where the proportionality factor *κ*(*n_sample_*, *λ*) is dependent on the refractive index *n_sample_* and the imaging wavelength *λ* as detailed in Supplementary Section S1. We used the silica data in Fig. S1**a** to determine *κ*(*n_silica_*, 440 *nm*) = 291 nm, and validated that the SP contrasts of both polystyrene (PS) and gold nanoparticles are consistent with this value when accounting for their respective refractive indices, see Fig. S1**d**, 1**e**. The silica proportionality factor *κ*(*n_silica_*, 440 *nm*) is used below to estimate EV sizes in SP.

### SPFI of liposomes with dual wavelength iSCAT imaging

Fig. 4 shows SPFI data on fluorescently stained liposomes which were prepared from DOPC lipid incorporating 1% DiI dye by extrusion through track-etched membrane filters with a 400 nm pore size (representative images in Fig. 4**a**). Extrusion using large pore sizes is known to produce liposomes with sizes both smaller and larger than the filter pore. This made the sample suitable to test SP of polydisperse samples, as the addition of the lipid dye allowed for an independent measure of total lipid content within each liposome. Fig. 4**c** shows SPFI scatterplots that were generated from the image data under 440 nm iSCAT illumination, and third root SP contrasts and DiI intensities were found to correlate well as shown by a linear fit to the data above 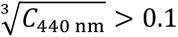 (dashed line). For many bright liposomes 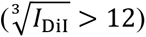, SP measurements underestimated liposome sizes due to the contrast inversion and many spots landed well above the fit line. Intriguingly, we found a maximum 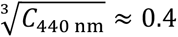 which is even larger than the 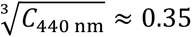 that we measured for 100 nm silica beads. This indicated that liposomes flattened upon adsorption on the glass surface which would result in a decrease of the interferometric phase factors and thus a delay of the contrast inversion. Flattening has been documented, and moreover liposomes are often used to form supported lipid bilayers^33^. We performed COMSOL simulations in Fig. 4**b** showing that flattening of solid particles while maintaining the total mass indeed results in a delay of the iSCAT contrast inversion without affecting the initial *d*^3^ scaling.

**Figure 4.**
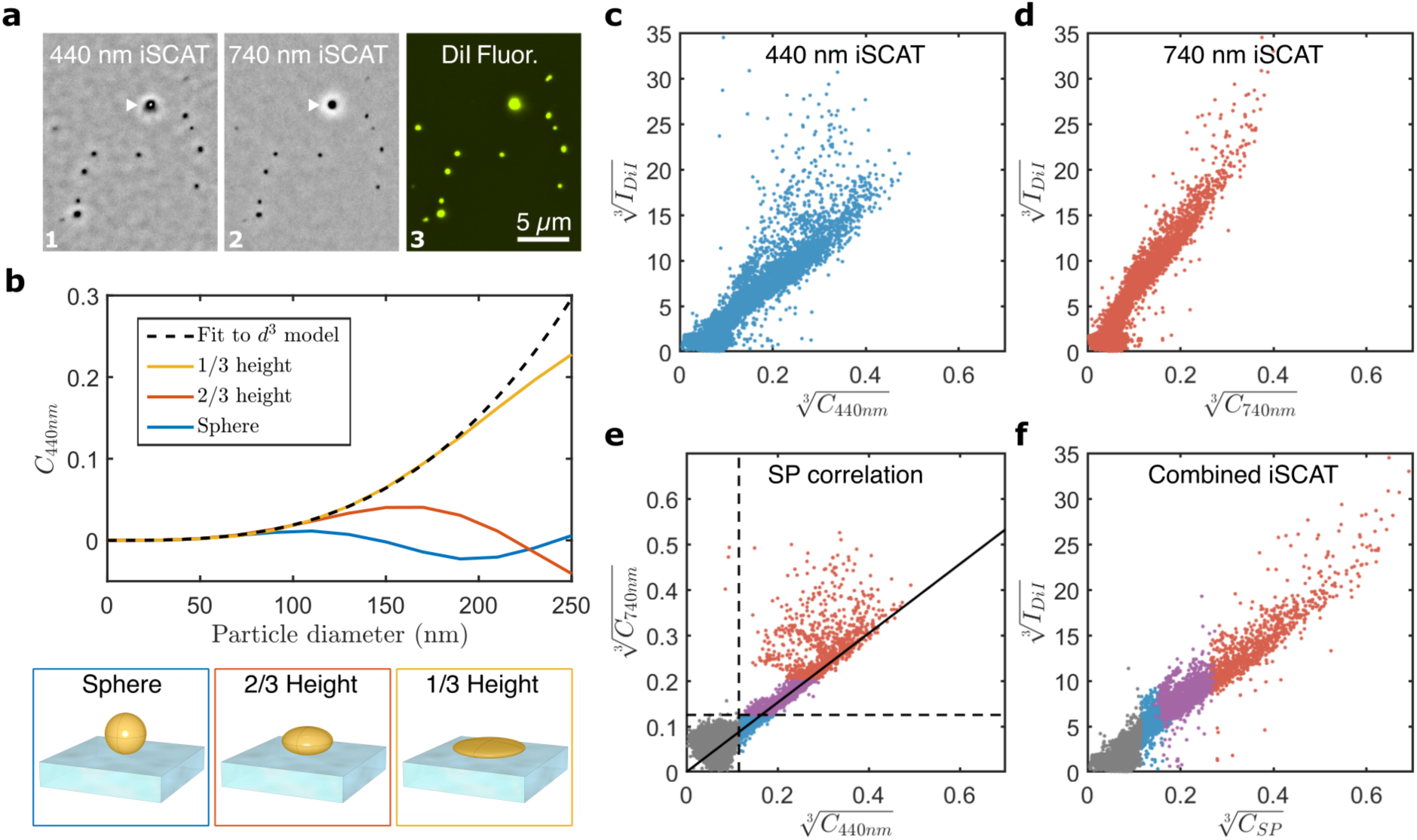
iSCAT at multiple wavelengths and fluorescence imaging of fluorescently stained liposomes. **a** Representative SP images of liposomes imaged at 440 nm (1, note white center in large dark spot indicated by a white arrowhead) and 740 nm (2) with aligned color scales (-1% to +0.5 %) and in fluorescence (3) to visualize their DiI stain. **b** FEM simulation of iSCAT contrasts at 440 nm of solid particles that remain spherical or flatten to 2/3 or 1/3 of the original height while maintaining a constant volume. Flattening allows larger particles to be measured by suppressing contrast inversion while increasing the maximal contrast. **c** SPFI scatterplot of DiI stained liposomes imaged with a 440 nm and **d** 740 nm iSCAT illumination, both showing a linear relationship albeit with different sensitivity (slope). **e** Scatter plot correlating the SP contrast at the two wavelengths. The purple dots indicate the size range at which the two contrast values are proportional to each other and is used to determine the proportionality factor η by a linear fit (solid black line). **f** Sizing of liposomes using both wavelengths, with blue dots measured at 440 nm, red dots measured at 740 nm and purple dots averaged between the two measurements using their proportionality factor η. Gray dots fall below the iSCAT detection threshold using either of the wavelengths.

We confirmed that many larger liposomes that underwent the contrast switch at 440 nm maintained dark iSCAT contrast when imaged under a longer 740 nm wavelength, as shown in Fig. 4**d**. Expectedly, this came at the cost of reduced sensitivity overall which we visualized by a SP scatterplot of both contrast measurements in Fig 4**e**. We found that in the region above the noise floor of the 740 nm channel and below the contrast inversion point of the 440 nm channel (purple dots in Fig. 4**e**) both measurements are roughly proportional to each other. Using a linear fit to the data (forcing a zero *y*-intercept), we determined a contrast proportionality factor *η* = 2.6 between the two wavelengths which was reasonably close to the theoretically predicted value of 2.8 based on wavelength scaling. Both measurements were combined to 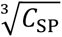 in Fig. 4**f**, where 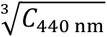 was used for small liposomes (blue dots), 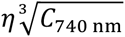 for large ones (red dots), and intermediately sized ones were averaged between the two measurements (purple spots).

Combining measurements in two wavelengths using a SP correlation plot (Fig. 4**e**) can thus maximize the dynamic range for sizing polydisperse samples at the expense of dual measurements.

### SPFI of extracellular vesicles

Small extracellular vesicles (sEV) from the supernatant of several cell lines were purified by size exclusion chromatography (SEC) and imaged using combined 440 nm and 740 nm illumination wavelengths. EV sizing with SP faces two principal challenges: firstly, EVs are expected to have a size-dependent refractive index since the contributions of EV membrane and lumen to the overall refractive index change with physical EV size and secondly, there can be additional variation in EV refractive index due to varying amounts of protein and nucleic acid loading. We addressed the first challenge using numerical calculations (Fig. S2**a**) and FEM simulations (Fig. S7), where we compared EVs modeled with solid sphere equivalent refractive index to core-shell models^25,34^ in which membrane and lumen contribute independently to EV scattering signals.

Both showed that within our expected sizing range, core-shell geometries lead to only minor corrections of EV sizes compared to the uniform models, which led us to adopt a single valued *n*_EV_ = 1.40 in our size calibration for simplicity. For completeness, all our plots in the following contain two *x*-axes indicating both raw and calibrated contrasts. The EV size calibration was done with 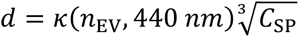, where the proportionality factor *κ*(*n*_EV_, 440 *nm*) = 369 nm was calculated from *κ*(*n*_silica_, 440 *nm*) as described in supplementary Section S1. With the detection limit 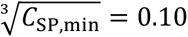 (Fig. 2**d**), this calibration results in an LOD for detecting EVs with 0.10*κ*_EV_ ≈ 37 nm diameters using our setup. Additionally, we found that experimentally measured SP contrasts at 440 nm illumination before contrast inversion can exceed maximum contrasts predicted by FEM simulations for both solid and core-shell EV models, suggesting that EVs, just as liposomes, can flatten upon surface adsorption^35^.

To address the challenge of random variations of EV refractive indices, SP was benchmarked against fluorescent stains which are commonly used for EV sizing. Fig. 5**a** shows an SPFI scatterplot of EVs from HT29 cells which were biotinylated via their membrane proteins using the membrane impermeable Sulfo-NHS-Biotin and then incubated with Alexafluor 674 (AF647)- labelled streptavidin. EVs were immobilized on glass coverslips pre-coated with poly-L-lysine (PLL) which is positively charged at neutral pH and captures EVs by electrostatic interaction.

**Figure 5.**
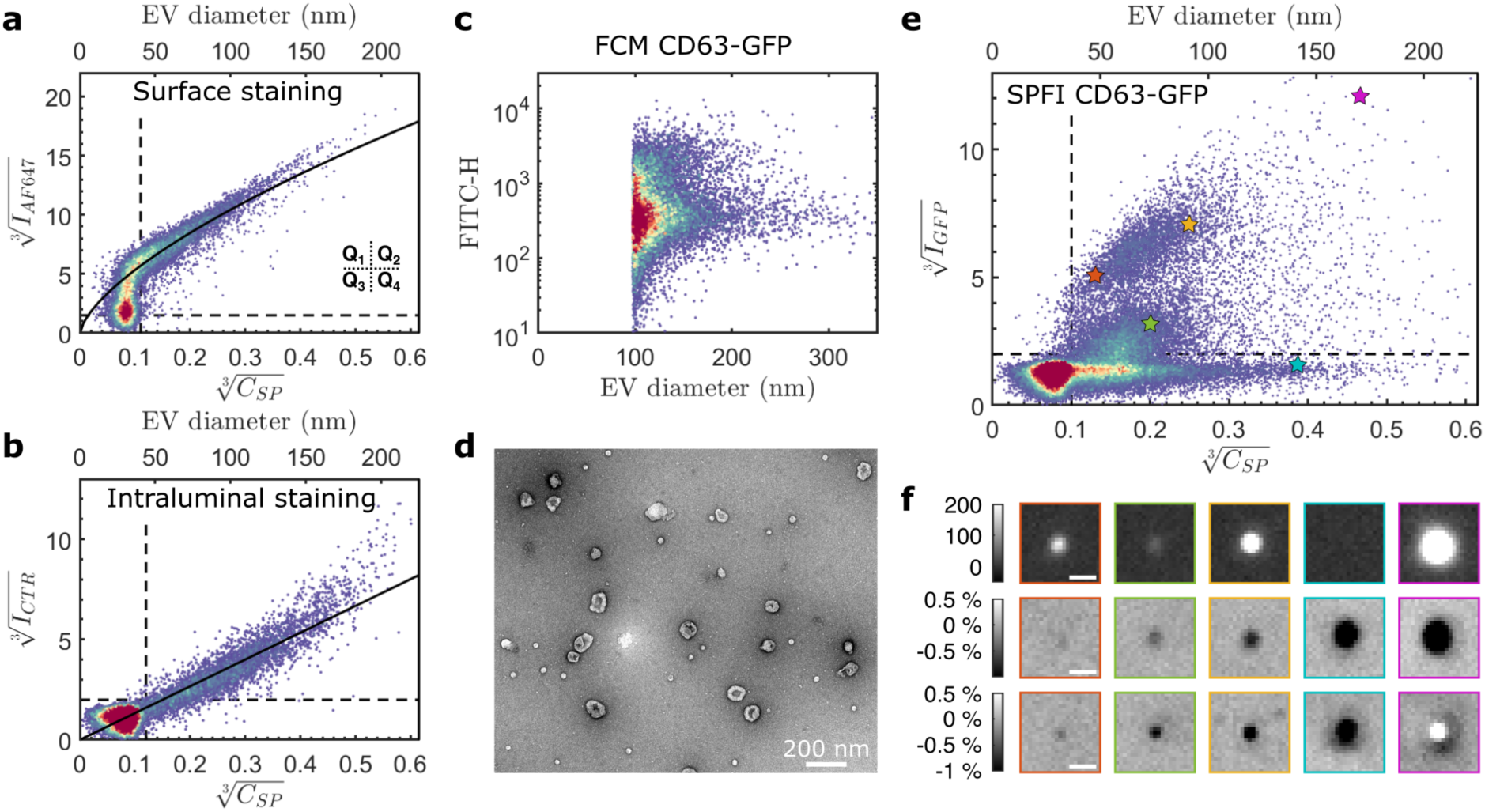
SPFI scatterplots correlating EV size with fluorescence intensity. **a** SPFI scatterplot of biotinylated EVs that were stained with Streptavidin-AF647. Dashed lines are contrast and intensity thresholds separating real detection events from image noise. Quadrants Q1 to Q4 apply to all scatterplots. The solid line is a model assuming homogenous labelling of the EV membrane proteins. **b** SPFI scatterplots of EVs stained using cell tracker deep red (CTR) where the solid line is a model assuming dye uptake in the lumen of the EVs. **c** FCM measurements of EVs from A431 cells expressing CD63-GFP measured in the fluorescein (FITC) channel. **d** Negative stain TEM image of the CD63-GFP EVs. **e** SPFI scatterplot of the CD63-GFP EVs showing two distinct clusters of GFP+ spots, one group with 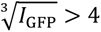 and EV sizes up to ∼100 nm and a smaller group with 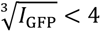 on EVs up to only ∼80 nm, either of which could not be resolved in FCM (see **c**). **f** Post-incubation images of selected EVs identified by star symbols in **f**. Top row: GFP fluorescence; middle row: 740 nm iSCAT; bottom row: 440 nm iSCAT. Panels can be matched to the spots in **f** by the colors of their borders. Intensity bars show raw iSCAT and fluorescence contrasts and scale bars are 500 nm.

The scatterplot was divided into four quadrants (Q1 to Q4, labelled in Fig. 5**a**) by applying SP contrast and fluorescence intensity thresholds found in control experiments (Fig. S9). We note that biotinylation is not specific to EVs and spots in Q1 on the top/left include many biotinylated soluble proteins as well as streptavidin molecules that were not washed off, however, by correlating the fluorescence data to label-free iSCAT, these spots can be efficiently removed from the data prior to further analysis. Assuming a constant membrane protein density for EVs of different sizes, the fluorescence signal intensity is expected to correlate with EV surface area. We tested this prediction by a fit to 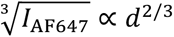 for 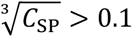, shown as solid line, which resulted in a correlation coefficient of *R*^2^ = 0.75. Any linear fit to the data would result in non-negligible *y*-axis intercepts which cannot be explained purely by detection sensitivities. Supplementary Fig. S9**d** shows additional data with membrane staining using MemBrite which shows a similar contrast behavior.

To further corroborate these finding, a second sample of HT29 derived EVs was stained with Cell Tracker Deep Red (CTR) and captured on a plasma cleaned glass surface, Fig. 5**b**. Since CTR accumulates in the EV lumen, the fluorescence intensity is expected to scale with the EV diameter and hence 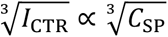. This scaling behavior was confirmed using a linear fit (solid line in Fig. 5**b**) to the datapoints above 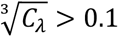 which resulted in a correlation coefficient *R*^2^ = 0.82. Similar to Fig. 5**a**, spots above 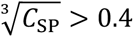 showed systematic deviation from the fit, indicating potential underestimation of EV sizes by SP due to limited sizing range at 740 nm (see also Fig. S10 for SPFI plots of the individual wavelengths). Fig. S11 shows additional experiments where EVs were co-stained with CTR and Carboxyfluorescein succinimidyl ester (CFSE) which both showed similar correlation with third root SP contrasts and similar detection sensitivities. Figs. 5**a** and 5**b** together with Figs. S9, S10, and S11 thus show that SP contrasts correlate well with four different fluorescent staining methods and can distinguish surface from bulk labelling on HT29 EVs.

Next, we highlight the utility of SPFI by comparing it to other single EV methods on EVs from A431 cells expressing CD63 fused to green fluorescent protein (CD63-GFP) in addition to their wildtype CD63 (Figs. 5**c**-5**g**). CD63 is a transmembrane protein from the tetraspanin family and a common marker of small EVs due to its high abundance and role in exosome biogenesis. Fig. 5**c** shows FCM data of this sample which had a LOD of *d* ≈ 100 nm and includes considerable technical noise as seen from the buffer control (Fig. S12, size calibration in Fig. S14). Fig. 5**d** shows a negative stain TEM image which confirmed the presence of many EVs of 50 nm and below. Fig. 5**e** shows an SPFI scatterplot of the data which detected EVs down to ∼37 nm in our calibration and shows non-uniform GFP fluorescence with two clear clusters. The first cluster comprises bright GFP^+^ spots 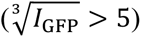 on EVs with diameters up to ∼100 nm with linear correlation between contrast and fluorescence. The second cluster consists of a much larger number of dim GFP^+^ spots 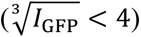 on EVs up to only ∼80 nm (see Fig. 5**f** for images of the EVs highlighted with arrowheads in Fig. 5**e**). The SPFI plot visually resolves the two clusters by combining label-free SP contrasts and CD63 expression data and the mismatch in size range between the clusters suggests that they represent different subpopulations. The same measurement in FCM, due to its higher detection threshold, could not resolve the two clusters. The iSCAT data furthermore allows to count the total EV number and we found that 8120 out of 17640 spots above the gating thresholds were positive for GFP, or in other words the overall EV labeling efficiency of the CD63-GFP transfected cells within our iSCAT detection range was 46 %.

Fig2. S12**a**, 12**b** show a comparison of the same sample captured on a plasma activated glass and on PLL coated glass, and we found that the overall shape of the SPFI plot was largely comparable and both GFP+ clusters are readily identifiable on either surface. SP contrast measurements differed minorly, which will be further investigated in future work to improve sizing and include capture bias of different surfaces. Fig. S13 furthermore shows the same EVs purified via a 10,000 g ultracentrifugation protocol which led to the inclusion of much larger EVs in the sample. We found that our current single pixel contrast measurements indeed limit SP sizing at 740 nm to EVs of ca. 200 nm. However, the data is suggestive that improved sizing including a full interferometric PSF model will increase the range in the future (see Fig. S13**d**).

## DISCUSSION

We described SP of EVs and nanoparticles using a common epifluorescence imaging setup, including the microscope with non-imaging based autofocus, LED-based illumination, and sCMOS camera, and an additional broadband 50:50 beamsplitter available for ∼ US$ 150. In our hands, SP was able to detect and size immobilized EVs down to *d* ≈ 37 nm. SP exceeds the detection sensitivities of many commercial FCM and NTA systems with a high throughput and high particle numbers by imaging over multiple FOVs, Fig. 6. The SP setup itself is similar to iNTA^25,26^ and in-solution measurements of absolute EV concentrations and hydrodynamic diameters are possible but will require more powerful illumination and high speed cameras to achieve similar LODs to SP. We propose the use of SP with fluorescence imaging as in SPFI as the main application which replicates the defining features of flow cytometry that can simultaneously capture fluorescence and label-free side scattering data. SP likewise complements EV fluorescence microscopy data with label-free nanoparticle enumeration and sizing, helps the identification of individual EVs and discriminate against unbound labels or substrate autofluorescence. Thanks to pre- and post-incubation imaging of immobilized EVs, high SP sensitivities which can be improved by longer averaging with common standard microscopy hardware, and in our hands performance was ultimately limited by the mechanical stability and focus drift of the setup (Fig. 2). Our setup makes no compromise on fluorescence imaging sensitivity, and we note that our SPFI workflow is compatible with super-resolution imaging^6^, EV subpopulation enrichment via affinity capture surfaces^6^, and cyclic immunofluorescence imaging^14^.

**Figure 6.**
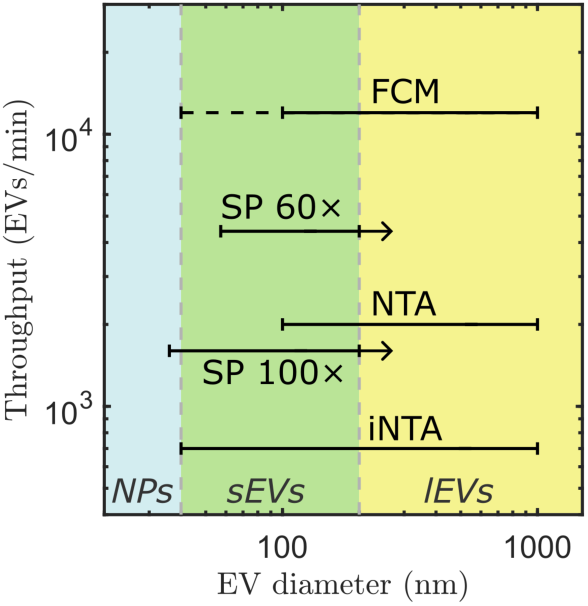
Comparison of SP against other label-free detection techniques. SP sizing range and throughput are plotted for imaging with a 100× as well as a 60× objective (see Fig. S15). The SP throughput is calculated by assuming a capture density of 400 EVs / (100 μm)^%^ together with the imaging parameters used in this manuscript (see methods, Fig. S5). FCM and NTA lines represent the commercial instruments used in this manuscript, whereas the dashed FCM line indicates the highest reported sensitivity on any FCM to our knowledge^9^. Note that NTA and iNTA^25^ report hydrodynamic diameters which are different to the physical EV size reported by SP and FCM. Shaded areas indicate commonly assumed size ranges for large EVs (lEVs), small EVs (sEVs) and smaller, non-membranous nanoparticles (NPs).

We present a SPFI pipeline which includes automated image pre-processing including iSCAT flatfield corrections and aberration corrections, FOV alignment, candidate spot detection, and contrast measurements using two SP illumination wavelengths. Single color SP pre- and post-incubation imaging of more than 10,000 EVs and nanoparticles over 20 FOVs could be completed within ∼7 min (more FOVs can be acquired if desired), consuming only 10 µl of sample per flow cell.

To establish label-free nanoparticle sizing with SP, we first imaged silica particles of known sizes and confirmed that SP contrasts *C* scale approximately with *d*^3^ for diameters. We found an upper sizing range for silica of about ∼90 nm when imaged under 440 nm wavelength, and about ∼150 nm diameter when imaged under longer 740 nm wavelength illumination, beyond which SP contrasts invert. We then compared SP contrasts of the 50 nm silica beads to 50 nm PS and gold nanoparticles and validated that the relative SP contrasts matched iSCAT theory when accounting for the difference in refractive indices. These results established a basis for sizing other nanoparticles and materials of known refractive index using the *d*^3^ scaling. A particular challenge to sizing EVs is that their refractive index can vary from the random incorporation of proteins and nucleic acids, potentially interfering with the expected size-dependency of contrasts in SP measurements. We used fluorescent dyes as independent measurements of cell culture derived EV sizes and found correlation coefficients between fluorescence intensities and SP contrasts of *R*^2^ = 0.75 for surface staining via biotinylation and *R*^2^ = 0.82 for intraluminal staining with CTR. This high correlation suggests that the *d*^3^ size dependence of SP contrasts can be strong enough to overwhelm refractive index variations and that SP can thus serve as a label-free EV sizing method. Given the large variety of possible EV samples, future studies are needed to establish the generality of this correlation, and we propose that fluorescent staining experiments are a practical way to verify SP for new sample types. Notably, SPFI was able to distinguish surface and volumetric stains via the relationship between size and fluorescence intensity *I* that was proportional to *d*^2^ and *d*^3^, respectively, which we believe to be the first direct experimental demonstration and further validation. We propose that size calibration using beads of various materials allows to standardize SP measurements between various microscopy setups, similarly to how size calibration is meant to standardize FCM measurements^8^. We note that the average EV refractive index is expected to vary as function of size (volume) because the proportion of lipid (surface based) changes. We thus performed FEM simulations of core-shell EV models to investigate how EV membrane and lumen with different refractive affect the size calibration. While EV refractive indices drop with increasing EV size, the simulations suggested that a solid sphere equivalent refractive index of *n*_EV_ = 1.40 is a good approximation to the average EV refractive index within our current SP sizing range. We propose that sizing accuracy and the upper sizing limit can likely be increased through improved image analysis in future work, either by explicitly modelling the full interferometric point spread function or by machine learning approaches. SPFI might also be used to further investigate EV refractive index variations and size dependence, for example by changing the refractive index of the surrounding media. We finally demonstrated the utility of SPFI for EV subpopulation discovery on CD63-GFP tagged EVs by comparing it to FCM. SPFI scatterplots revealed that GFP was present in EVs below 100 nm in size in two distinct clusters that differed in both GFP brightness and SP contrasts, whereas FCM only picked up diffuse expression on EVs larger than 100 nm.

A corollary of the above discussion is that the capture conditions including buffer, pH, salt concentration, and surface and EV charge could influence EV physisorption and introduce capture or sizing bias. We mainly captured EVs by electrostatic interaction using positively charged PLL coated coverslips (Supplementary Video 1). We found, however, that EVs could also be immobilized on negatively charged surfaces in PBS buffer, suggesting screening of the EV charge and major contribution by Van der Waals forces to EV adsorption (Supplementary Video 2). We found that electrostatic attraction on PLL increased capture rates but could also increase EV flattening on the surface. Flattening without material loss results in a reduction of the interferometric phase resulting from the out-of-plane material. This extends the range in which *C* ∝ *d*^3^ as it delays the contrast inversion, which could increase the range of SP. Strong deformations, however, might lead to fluid loss from the EV lumen and thus additional complications for EV sizing. Future work will investigate whether EV deformability in different capture environments could open the door to all optical single vesicle stiffness or density measurements using SP.

## Conclusions

SPFI of EVs is readily accessible by the addition of a 50: 50 beam splitter to any modern epifluorescence microscope and runs on an openly available image processing pipeline (https://github.com/junckerlab/SPFI). SP on pre- and post-incubation images can detect and size EVs from *d* ≈ 37 nm to > 200 nm, and its combination with fluorescence imaging generates rich multimodal single EV datasets that could help identify EV characteristics and clusters in > 500 EVs per FOV, and > 10,000 in each microfluidic flow cell in our experiments. SPFI can distinguish membrane from intraluminal dyes in EVs, captures deformation of EVs, and can reveal subpopulations that are only distinguishable in EVs with *d* < 100 nm. Finally, SPFI should be broadly applicable to EVs and other rigid or soft, synthetic or biological, nanoparticles.

## Methods

### Microscopy setup

The iSCAT system was built on an inverted fluorescence microscope (Nikon TI2) using a 100× plan apochromat (λ series, Nikon) of NA 1.45 with a 1.5× intermediate magnification lens (inbuilt feature of the Ti2). We performed additional characterization of SP detection sensitivity using a 60× objective shown in Fig. S14, however, due to reduced sensitivity all experimental data shown was taken with the 100× objective. Imaging was done on a Prime95B (Photometrics) which has a full frame 118×118 µm^2^ FOV size with 1608×1608 pixel sensor, however, measurements were taken with a reduced 1200×1200 pixel region of interest (ROI) to ensure optimal focus throughout the ROI. Illumination was provided by a Spectrapad (Lumencor) via a standard fixed large FOV epifluorescence illuminator (Ti2-LAPP *fixed main branch*, Nikon) with built-in fly-eye lens. The illuminator has an adjustable aperture and two alignment screws which both affected the achievable iSCAT contrasts (data not shown). Aperture alignment and opening was optimized for highest contrast of 50 nm silica particles before taking all data in this manuscript. iSCAT using multiple FOVs furthermore relied on an LED based reflectance (non-imaging) autofocus (*Perfect Focusing System*, Nikon).

The fluorescence measurements were taken using a pentapass dichroic mirror (FF409/493/573/652/759-SP01, Semrock) and pentapass emission filter (FF01-432/515/595/681/809, Semrock). The emission was additionally filtered using 515/30 nm (GFP), 595/40 nm (SPI) or 680/42 nm (AF647) filter sets (Semrock) in an automated filter turret in front of the camera. Maximum illumination power of the Spectrapad was used in all channels and images were averaged 8× with 100 ms exposure time each.

iSCAT imaging used the same light source (Spectra X Light Engine, Lumencor) which provides 370 mW output power in the 440/20 nm channel and 80 mW in the 740/20 nm channel at the maximum power setting according to manufacturer specifications. A filter cube with a widefield 50:50 beam splitter (BSW10R, Thorlabs) was added in the filter turret to switch between fluorescence and iSCAT modes. The frame exposure was maximised within the dynamic range of the camera, which resulted in exposure times around 4 ms at 440 nm and 25 ms at 740 nm.

This allowed to image ROI of 512×512 pixels at 95 FPS, or 1200×1200 pixels at 40 FPS.

#### iSCAT noise characterization

The noise characterization in Fig. 2 was performed using the same settings as for regular imaging in FOVs of 512×512 pixels with camera exposure of 4.8 ms. The expected noise was calculated using the camera bit depth of *δ* = 2^16^, well depth of *γ* = 80,000 e-, quantum efficiency of *η* = 95 % and readout noise *v* = 1.6 *e*^-^. Due to illumination inhomogeneities, the maximal exposure is only achieved in a small region in the center of each frame while it drops off substantially towards the edges. We adjusted camera exposure time to achieve highest single pixel exposure of 98.5 % saturation which resulted in an average pixel exposure of *ρ* = 75.4 %. This corresponds to 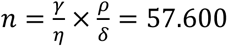 captured photons per pixel for single frame averaging and allows to calculate the photon shot noise as *σ* = √*n*, shown as solid line in Fig. 2**c**. Since the camera readout noise was limiting the dynamic range to 50,000, we also calculated an adjusted noise floor by accounting for the readout noise as *σ*′ = *σ*_sn_/√υ.

All points in Fig. 2**c** were generated using a dataset containing *N* = 5000 images (taken at a frame rate of ∼500 images/s) of an empty coverslip with an average pixel saturation of ∼90 % (similar to Fig. 2**a**). For each amount of averaging, we computed two images, *I*_U_ and *I*_E_, where *I*_U_ is the average of the first *n* uneven frames, and *I*_E_ is the average of the first *n* even frames of the original time series. This interleaved averaging of the frame made the analysis robust against slow illumination power fluctuations. We then calculated the difference *I*_diff_ = *I*_E_ − *I*_V_to eliminate all spatial information of the individual frames such as illumination inhomogeneities. We calculated the noise in *I*_diff_ using the standard deviation *σ*_diff_. Since both individual images *I*_E_ and *I*_U_contribute to *σ*_diff_in equal parts, we can finally estimate the image noise for an iSCAT image averaging 2*n* frames as *σ_n_* = 1/√2 *σ*_diff_.

To estimate the image noise after flatfield correcting the raw iSCAT frames, we generated two extra images, *FF*_U_ and *FF*_E_, to mimic the real flatfield images. The stand-in flatfield images were summed from *n*_FF_ = {1, 10, 100, 1000} additional frames of the timeseries which were not already used to generate *F*_E_ and *F*_U_. *F*_U_ and *F*_E_ were then divided and subtracted as 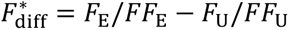 such that the resulting noise can be calculated via the standard deviation of 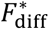.

We note that additional background was present in real experiments which is described in Supplementary Section S6.

#### Flow cell preparation and sample incubation

All flowcells used in this manuscript used #1.5 glass coverslips (Fisher Scientific) as substrate. Coverslips were sonicated in acetone and isopropyl alcohol for 5 min each and rinsed in distilled water before blow-drying with nitrogen. The coverslips were plasma treated for 1 min and sealed into a custom hybriwell chamber (Grace bio-labs). The flow cells have a volume of 10 µl with a rectangular center area of 6 mm × 8 mm with conical taper regions at the short sides (15 deg half angle) which leads to 1.5 mm diameter in- and outlets. The height of the flow cells is 150 µm and a total of five such flow cells are contained on each hybriwell chamber. Initial filling of the flow cells is aided by capillary pressure after manual pipetting into the flow cell inlet. Subsequent sample filling or washing of the flow cell was done by pipetting onto the inlet while gently pressing a folded lint-free tissue (Kimwipe, Kimberly-Clark Professional) to the outlet. Flow cells were filled with the respective buffer (see sample descriptions) before taking the pre-incubation images. 10 µl of sample was flushed into the flow cell and EV capture was monitored using iSCAT imaging (without flatfield processing). After the appropriate density was achieved (typically 1 – 3 mins), the flow cell was flushed with 100 µl of buffer and the post-incubation images were taken.

#### Calibration samples

Aminated silica nanoparticles with diameters of 22 nm, 52 nm, 77 nm, 97 nm, and 208 nm (SIAN20-25M, SIAN50-25M, SIAN80-25M, SIAN100-25M, SIAN200-25M) were purchased from nanocomposix (San Diego, USA). Prior to use, the particles were sonicated and then diluted to approximate concentrations of 1 × 10^9^ particles per ml in 2 mM acetate buffer at a pH of 5 where they attained positive surface charge.

Citrated gold nanoparticles with diameters of 10 nm, 20 nm, and 50 nm (A11-10-CIT-DIH-1-1, A11-20-CIT-DIH-1-1, A11-50-CIT-DIH-1-1) were purchased from Nanopartz Inc. (Loveland, USA) and diluted in milliQ water before use.

NIST traceable Nanosphere PS Size Standards with diameters of 51 nm, 81 nm, and 100 nm (2050A, 3080A, 3100A) were obtained from Thermo Fisher Scientific (Mississauga, Canada) and diluted in milliQ water before use.

Liposomes were prepared from DOPC lipid (Avanti lipids, Alabaster, USA, cat. no. 850375) in chloroform solution which was spiked at a 1 % molar ratio with DiI dye in ethanol. The solvents were evaporated in a test tube under vacuum overnight. The lipid film was resuspended in PBS to a molar concentration of 10 mM and allowed to swell and form large multilamellar bubbles for 30 mins. The solution was then extruded by passing 21 times through a filter with pore size of 400 nm (Avanti lipids) and diluted further for the incubation.

#### Cell culture and EV isolation

Human colorectal cancer cells HT-29 (ATCC) and CD63-GFP modified A431 cells (CD63 with C-terminal fusion of GFP, a gift from the lab of Janusz Rak^36^) were cultured in Dulbecco’s modified Eagle Medium (DMEM) culture medium with 10 % FBS and 1 % (vol/vol) penicillin-streptomycin. The cells were incubated in a humidified incubator at 37 °C and 5 % CO_2_and grown to about 60 % confluency. The medium was then replaced by DMEM with 5 % EV-depleted FBS and the cell supernatant containing the EVs was harvested after ∼48 hrs. The supernatant was spun at 400 g for 15 min to pellet cell debris and filtered by 450 nm syringe filters to remove larger EVs. For each EV sample, 5 ml supernatant was concentrated to 500 µl using 100 kDa Amicon Ultra-0.5 centrifugal filters (Millipore) for 5 mins at 5,000 g. The concentrated supernatant was purified using single use qEV.70nm size exclusion columns (IZON) with 0.8 ml void volume after sample loading and 0.6 ml sample collection volume.

#### CTR staining of EVs

EV samples were stained twice with CTR after it was found that the correlation of size to fluorescence intensity in the SPFI plots could be considerably increased compared to single rounds of staining (data not shown). Single staining resulted similar high staining efficiencies as reported for double staining (i.e. the number CTR+ EVs) but many large EVs were only relatively dim and variations between EVs of the same SP contrast were large, whereas SP size distributions were comparable regardless of staining. For the first staining round, 2 µl of Cell Tracker Deep Red dye (Invitrogen) at 1 mM concentration was added to 500 µl concentrated cell supernatant and incubated for 2 hrs at 37 °C. EVs were then purified using IZON qEV.70nm size exclusion columns as described above. Afterwards, a second round of staining was done using 2 µl of CTR in 500 µl purified EVs for 2 hrs at 37 °C. Excess dye was removed in six washes with PBS using 100 kDa Amicon Ultra-0.5 centrifugal filters (Millipore) for 5 mins at 5,000 g (each spin concentrates 500 µl sample to ∼15 µl).

#### Biotinylation and AF647-Streptavidin staining of EVs

Biotinylation was done using the EZ-Link Sulfo-NHS-LC-LC-Biotinylation Kit (Thermo Fisher Scientific, Mississauga, Canada). 0.5 mg of Sulfo-NHS-LC-LC-Biotin was added into 500 µl of purified EVs (typically ∼10^9^ EVs per ml measured by NTA) and incubated for 30 min at room temperature. Unreacted biotin was removed in six washes with PBS using 100 kDa Amicon Ultra-0.5 centrifugal filters (Millipore) for 5 min at 5,000 g. Subsequently, 1 µl of Alexa Fluor 647 labelled streptavidin at 1 µg/ml was added and incubated for 20 min. Excess streptavidin was removed in six washes with PBS using 100 kDa Amicon Ultra-0.5 centrifugal filters (Millipore) for 5 min at 5,000 g.

#### Flow cytometry measurements

All FCM data presented here was taken using a CytoFLEX-S Flow Cytometer (Beckman Coulter) equipped with 405 nm, 488 nm and 638 nm lasers, and operated using CytExpert Software. The 405 nm violet laser was used for side scattering (VSSC) and the 488 nm laser for forwards scattering (FSC). VSSC gain was set to 200, SSC gain to 500 and FITC gain to 1000, the VSSC threshold was set to 1400. Before each sample measurement, the system was washed using Azide buffer at pH 5 for silica nanoparticles, ultrapure water for PS calibration beads and PBS for EV samples. An event rate below 1000 per second was ensured for these buffer controls. Samples were loaded and after the event rate became stable and measurements were taken for 60 s each.

The size calibration (Fig. S13) was done in FCMPass^34^ v. 4.2.14 using 81 nm, 100 nm, 152 nm, 203 nm, 240 nm, 303 nm, 345 nm, 401 nm and 453 nm polystyrene beads (3080A, 3100A, 3150A, 3200A, 3240A, 330A, 3350A, 3400A, 3450A) purchased from Thermo Fisher Scientific (Mississauga, Canada). The calibration was done using the default *Average RI* core-shell model with a 5 nm thick shell of refractive index 1.4863 and core refractive index of 1.3859.

#### NTA measurements

NTA measurements were performed on a Nanosight (Malvern Panalytical, Malvern, UK) NTA 3.4 using a 488 nm laser. Five measurements of 60 s each were taken per sample at a syringe pump speed of 40 at 25 frames per second and detector threshold of 5. The camera level was set to 14 for the SEC sample and 10 for the 10,000 g UC sample to keep overexposure below 10 % of traces.

#### Negative stain TEM imaging

Copper grids were negatively charged for 20 sec at 20 mA and 5 μL of EV sample was incubated without dilution after SEC purification for 10 min. The grids were washed three times in ultrapure water for 30 s each and fixed in 2 % glutaraldehyde solution for 2 min followed by additional three washes in ultrapure water for 30 s each. Staining was performed with a 2% uranyl acetate solution for 45 sec, the grids were blotted dry using filter paper and residual moisture was left to evaporate for 1 hr. TEM images were taken on a FEI Tecnai 12 at a working voltage of 120 kV.

## Data availability

Matlab scripts for evaluating SPFI data, including all image pre-processing, alignment and spot localization algorithms detailed in Supplementary Sections 2, 3 and 4 are available via https://github.com/junckerlab/SPFI.

All raw data supporting the main findings of this manuscript can be found at https://doi.org/10.5281/zenodo.10208005.

## Acknowledgments

We thank Carl Fabian Svahn and Hugues Martin for useful discussions and comments on the manuscript. We gratefully acknowledge Hélène Pagé-Veillette at the McGill University Health Centre (MUHC) for assisting with the FCM measurements and providing polystyrene bead calibration data. We also thank Jennie Mui and Kelly Sears for assistance with TEM imaging at the Facility for Electron Microscope Research (FEMR) at McGill University. We are thankful for financial support from Genome Canada’s Disruptive Innovation in Genomics, the Natural Science and Engineering Research Council of Canada’s Discovery Grant (NSERC, RGPIN-2016-06723), and the Canadian Cancer Society’s Operating Grant 943871. D.J. acknowledges support from a Canada Research Chair. A.W. is supported by Schmidt Science Fellows, in partnership with the Rhodes Trust.

## Declaration of Interest Statement

The authors declare that the research was conducted in the absence of any commercial or financial relationships that could be construed as a potential conflict of interest.

## Notes

### Competing Interest Statement

The authors have declared no competing interest.

### Summary of Updates

Revision in response to reviewer comments notably including contextualization and generalization of SPFI and size photometry in particular and including new measurements to calibrate and validate sizing.

