## Supplementary Information for "Size photometry and fluorescence imaging of immobilized immersed extracellular vesicles"

### S1: iSCAT imaging theory

iSCAT has been developed as a label-free microscopy method for the detection of ultrasmall particles that are captured on a transparent substrate or float close to it[1]. It has gained popularity for optically weighing single molecules in a process called mass photometry[2,3] and it has occasionally been used on larger nanoparticles and EVs as well[4–6].

The following paragraph presents the theoretical background including the mathematical formulas used to calibrate the particle sizes from their contrasts following Ref. [7]. Small dielectric particles illuminated with optical power $P_{\mathrm{inc}}$ scatter light proportional to their scattering cross-section $\sigma$ as $P_{\mathrm{scat}} \propto \sigma P_{\mathrm{inc}}$. The scattering cross section is dependent on the illumination wavelength $\lambda$, refractive indices of both particle ($n_{\mathrm{EV}}$) and medium ($n_{\mathrm{m}}$), as well as the volume $V$ of the EV. It can be calculated as:

$$\sigma = \frac{8}{3}\pi^2|\alpha|^2(\lambda)^{-4} \qquad (1)$$

Here, $\alpha$ is the particle polarizability given by:

$$\alpha = 3V\left(\frac{{n_{\mathrm{EV}}}^2 - {n_{\mathrm{m}}}^2}{{n_{\mathrm{EV}}}^2 + 2{n_{\mathrm{m}}}^2}\right) \qquad (2)$$

The total scattered power hence scales as $P_{\mathrm{scat}} \propto \sigma \propto V^2 \propto d^6$ where $d$ is the particle diameter for spherical particles. This strong size dependence makes particle detection by pure scattering (i.e. darkfield microscopy) exceedingly challenging below a hundred nanometers or so since the scattered light signal is often below the camera readout noise or overwhelmed by stray light. Even if sufficient sensitivity is achieved, the strong diameter dependence practically limits the dynamic range of the measurement as larger particles saturate the detector.

iSCAT improves on pure scattering detection by interfering the scattered field $E_\mathrm{s}$ with a strong reference field $E_\mathrm{r}$. It can be conveniently implemented by imaging the particle perpendicular through a glass surface where the reference field is generated by the back-reflection of a small fraction of incoming light from the glass surface (see Fig. 1**a** of the main text). The concept of interferometric detection is commonly described by splitting the detected power $P_\mathrm{det}$ into the following contributions:

$$P_{det} = |E_\mathrm{r} + E_\mathrm{s}|^2$$

$$= |E_\mathrm{r}|^2 + |E_\mathrm{s}|^2 + 2|E_\mathrm{r}||E_\mathrm{s}| \cos\varphi$$

$$= P_\mathrm{ref} + P_\mathrm{scat} + P_\mathrm{int} \cos\varphi \qquad (3)$$

The first term $P_\mathrm{ref} = |E_\mathrm{r}|^2$ is the constant background from the back-reflection at the glass surface and $P_\mathrm{scat} = |E_\mathrm{s}|^2$ is the pure scattering from the nanoparticle. The cross-term $P_\mathrm{int} = 2\sqrt{P_\mathrm{scat}}\sqrt{P_\mathrm{ref}} \cos\varphi$ is the interference signal between the two. It scales as $P_\mathrm{int} \propto \sqrt{P_\mathrm{scat}} \propto d^3$ which is much more favorable that the previous $P_\mathrm{scat} \propto d^6$ of pure scattering detection and explains the high sensitivity of interferometric detection for small particles. The interference term in this simplified description contains an interferometric phase $\varphi$ which lumps together the path difference of reflected and scattered fields, the Gouy phase[8],-and contributions from the complex valued dielectric permittivity of the particle. For small particles $\cos\varphi \approx -1$ such that particles contrasts are darker than the image background. We define the SP contrast $C_\mathrm{SP} = \frac{P_\mathrm{det}}{P_\mathrm{ref}} - 1 \approx \frac{P_\mathrm{int}}{P_\mathrm{ref}}$ such that darker than background spots gain positive contrast values. This results in the scaling law $C_\mathrm{SP} \propto d^3 - O(d^6)$ used throughout the main text (referred to as $d^3$ law) which is valid in the limit of small particles compared to the imaging wavelength.

We calibrated EV sizes in Fig. 5 on the main text using $d = \kappa \sqrt[3]{C_{\mathrm{SP}}}$ where the proportionality factor $\kappa(n_{\mathrm{EV}}, \lambda)$ to third root contrasts $\sqrt[3]{C_{\mathrm{SP}}}$ is dependent on the refractive index $n_{\mathrm{EV}}$ and the imaging wavelength $\lambda$. It is possible to calculate $\kappa(n_{\mathrm{EV}}, \lambda)$ for EVs (or any other sample material of known refractive index) based on a reference, for example $\kappa(n_{\mathrm{silica}}, \lambda)$ by setting $\kappa( n_{\mathrm{EV}}, \lambda) \sqrt[3]{C_{\mathrm{SP}}(n_{\mathrm{EV}}, \lambda)} = \kappa(n_{\mathrm{silica}}, \lambda) \sqrt[3]{C_{\mathrm{SP}}(n_{\mathrm{silica}}, \lambda)}$ . While not strictly necessary, we performed our calibration for each imaging wavelength independently such that we can write $C_{\mathrm{SP}}(n_{\mathrm{EV}}) \propto \sqrt{P_{\mathrm{scat}}} \propto \alpha \propto \left|\frac{n_{\mathrm{EV}}{}^2 - n_{\mathrm{m}}{}^2}{n_{\mathrm{EV}}{}^2 + 2n_{\mathrm{m}}{}^2}\right|$ using equation 2, and analogously for $C_{\mathrm{SP}}(n_{\mathrm{silica}})$.

Overall, we thus get

$$\kappa(n_{\mathrm{EV}}) = \kappa(n_{\mathrm{silica}}) \sqrt[3]{\left|\frac{n_{\mathrm{silica}}{}^2 - n_{\mathrm{m}}{}^2}{n_{\mathrm{silica}}{}^2 + 2n_{\mathrm{m}}{}^2}\right| / \left|\frac{n_{\mathrm{EV}}{}^2 - n_{\mathrm{m}}{}^2}{n_{\mathrm{EV}}{}^2 + 2n_{\mathrm{m}}{}^2}\right|}. \quad (3)$$

The reference proportionality factors $\kappa(n_{\mathrm{silica}})$ can readily be calculated by measuring SP contrasts $C_{\mathrm{SP}}(n_{\mathrm{silica}})$ on a calibration dataset with silica beads of known sizes $d$ via $\kappa(n_{\mathrm{silica}}) = d/\sqrt[3]{C_{\mathrm{SP}}(n_{\mathrm{silica}})}$ .

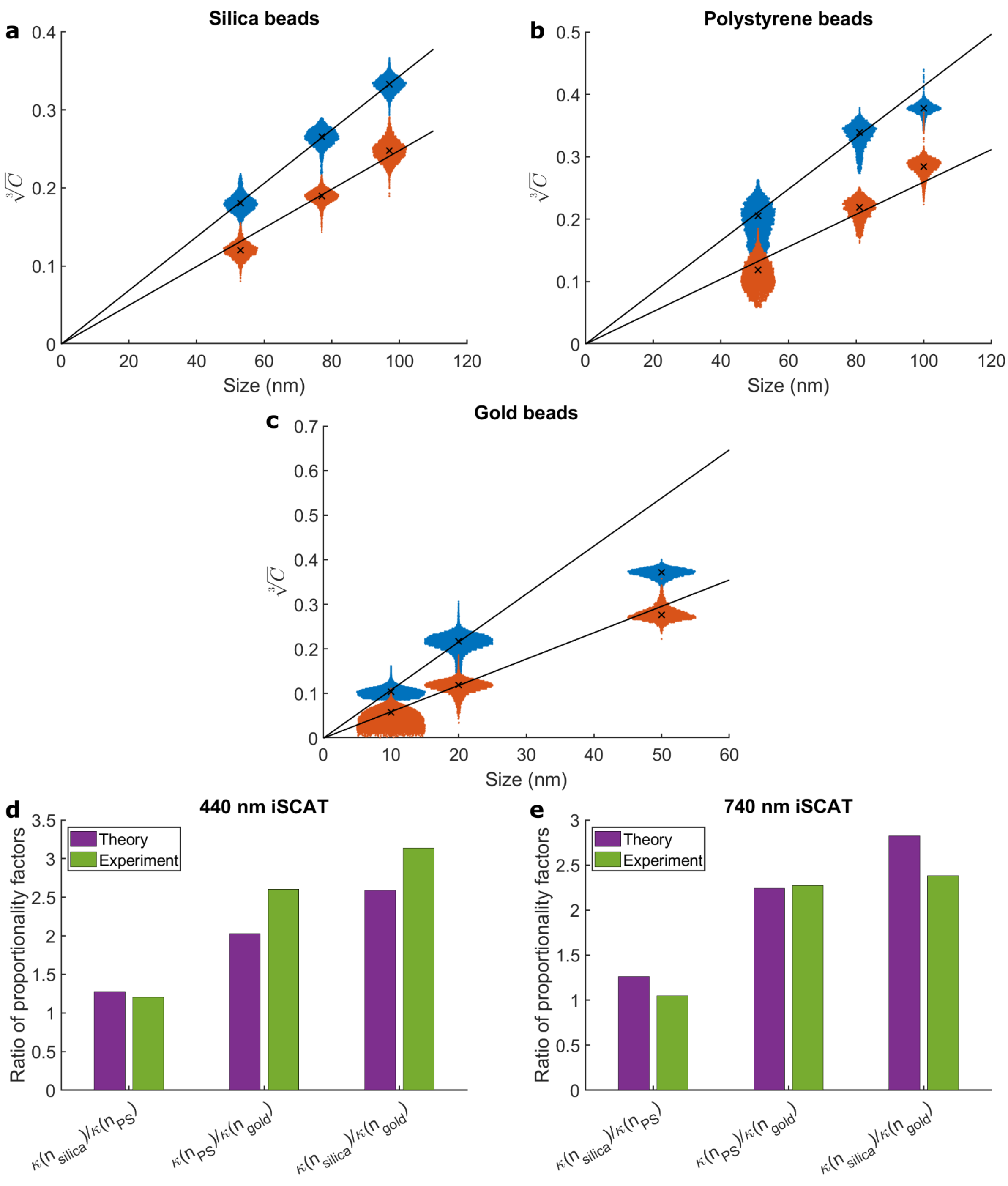

Figure S1: Calibration of SP contrasts using size standards of various materials. **a** Violin plots of SP contrasts of silica beads with nominal diameters 53 nm, 77 nm, and 97 nm imaged under 440 nm (blue) and 740 nm (red) illumination. Solid lines are linear fits using the median contrasts (crosses) of the 53 nm and 77 nm beads. **b** Violin plots of SP contrasts of polystyrene beads with nominal diameters 51 nm, 81 nm, and 100 nm. Solid lines are fits using the median contrasts of the 51 nm and 81 nm beads. **c** Violin plots of SP contrasts of gold beads with nominal diameters 10 nm, 20 nm, and 50 nm. Solid lines are fits using the median contrasts of the 10 nm and 20 nm beads. **d** Ratio of SP contrasts for the different

We show our calibration data for $\kappa(n_{\mathrm{silica}})$ together with validation on polystyrene (PS) and gold particles in Fig. S1. We note that for technical reasons, the sensitivity of our setup was slightly reduced during the preparation of this manuscript and that the silica data shown in Fig. 3 of the main text was taken on the older version of our setup. We kept Fig. 3 of the main text because it has more datapoints than Fig. S1, however, the revised silica dataset shown in Fig. S1 is the one used for EV size calibration because all other data in the manuscript was taken using the same setup and settings. Fig. S1**a** shows violin plots of $\sqrt[3]{C}$ for silica beads with nominal sizes 53 nm, 77 nm, and 97 nm. The contrasts distributions represent the main clusters found in SP plots (similar to Fig. 3**b**) and crosses mark the median contrast values. Solid lines are least square fits to all median values using $\sqrt[3]{C} = \eta d$ to calculate proportionality factors $\eta$, from which $\kappa(n_{\mathrm{silica}}) = 1/\eta$ were determined to be $\kappa(n_{\mathrm{silica}}, 440\ nm) = 291$ nm and $\kappa(n_{\mathrm{silica}}, 740\ nm) = 402$ nm. Fig. S1**b** shows a calibration dataset for polystyrene beads (PS) with nominal diameters 51 nm, 81 nm, and 100 nm from which we calculated $\kappa(n_{\mathrm{PS}}, 440\ nm) = 242$ nm and $\kappa(n_{\mathrm{PS}}, 740\ nm) = 385$ nm by fitting the 51 nm, 81 nm beads. Lastly, Fig. S1**c** shows a dataset with gold spheres of 10 nm, 20 nm, and 50 nm nominal size with which we calculated $\kappa(n_{\mathrm{gold}}, 440\ nm) = 93$ nm and $\kappa(n_{\mathrm{gold}}, 740\ nm) = 169$ nm by fitting the 10 nm and 20 nm spheres. We validated that the calibration factors are consistent with equation 3 in Figs. S1**d**, 1**e** by calculating a ratio of proportionality factors for all three material combinations. We compared the experimentally found ratios to theoretical ones that were calculated using bulk literature refractive indices of $n_{\mathrm{silica,440\ nm}} = 1.4663$ and $n_{\mathrm{silica,740\ nm}} = 1.4544$ for silica[9], $n_{\mathrm{PS,440\ nm}} =$

1.6161 and $n_{\mathrm{PS,740\ nm}} = 1.5808$ for PS[10], and $n_{\mathrm{gold,440\ nm}} = 1.4174 + 1.9322i$ and $n_{\mathrm{gold,740\ nm}} = 0.13689 + 4.4056i$ for gold[11] where the imaginary part describes gold absorption. We found that all ratios matched theoretical predictions for both wavelengths very well, in particular $\kappa(n_{\mathrm{silica}})/\kappa(n_{\mathrm{PS}})$ which validates our use of $\kappa(n_{\mathrm{silica}})$ for the EV calibration. We note that the theoretical predictions for SP contrasts of gold particles can likely be refined by adjusting the complex refractive index $n_{\mathrm{gold}}$ to account for size-dependent plasmon resonance effects.

Our EV calibration was done using equation 3 by assuming an average EV solid sphere equivalent refractive index of $n_{\mathrm{EV}} = 1.40$ which yields $\kappa(n_{\mathrm{EV}}, 440\ nm) = 369$ nm at 440 nm illumination (note that $\kappa(n_{\mathrm{EV}}, 440\ nm)$ is also used to calibrate two-color measurements). To decide on the value of $n_{\mathrm{EV}}$, we used core-shell models with different refractive indices for the core of $n_{\mathrm{core}} = 1.3658, 1.3859$ and 1.4060 which were taken from FCMPass v. 4.1.1. The shell thickness was $t = 5$ nm and the shell refractive indices $n_{\mathrm{shell}} = 1.48$. For any EV diameter $d$, the solid-sphere equivalent refractive index can be calculated via $n_{\mathrm{solid}}(d) = \frac{V_{\mathrm{shell}} n_{\mathrm{shell}} + V_{\mathrm{core}} n_{\mathrm{core}}}{V_{\mathrm{shell}} + V_{\mathrm{core}}}$ where $V_{\mathrm{shell}}$ and $V_{\mathrm{core}}$ are shell and core volumes, respectively. We performed numerical calculations to calibrate EV sizes in the core-shell model by estimating an initial size $d$ using $n_{\mathrm{solid}} = 1.40$ in the formulas above, then updating $n_{\mathrm{solid}}(d)$ with the just found estimate of $d$, and iterating these two steps until $d$ converged to within 0.1 nm between iterations. We show the results of these calculations in Fig. S2**a**, indicating that $n_{\mathrm{EV}} = 1.40$ is a good estimate for all three core-shell models within our sizing range. While the core-shell calibration would improve sizing accuracy slightly, we opted to use the single solid sphere refractive index calibration throughout the manuscript since it allows to overlay raw third root contrast values $\sqrt[3]{C_{\mathrm{SP}}}$ with the calibration data without distorting axes. Fig. S2**b** shows the variation in the setup calibration $\kappa_{\mathrm{EV}}$ depending on the refractive index values for both silica and EVs that are used.

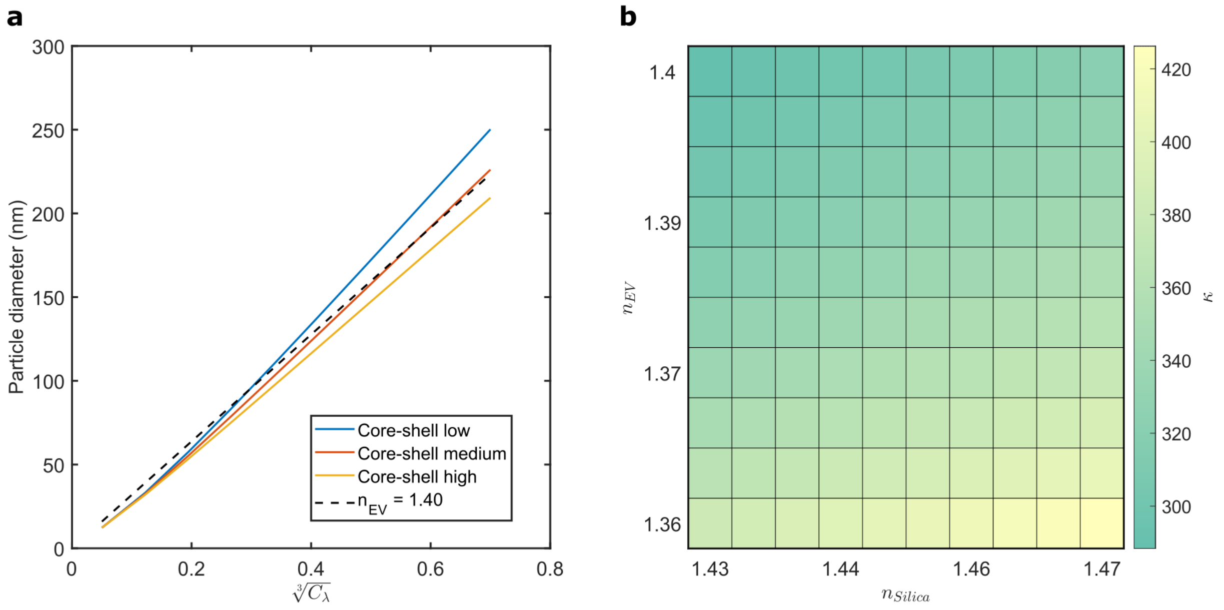

Figure S2: EV size calibration and dependence on the refractive index. **a** EV size calibration using a solid sphere equivalent refractive index model compared to calibration curves for three different core-shell models. Low, medium and high refractive index core shell models use core refractive indices of $n_{\mathrm{core}}$ =1.3658, 1.3859 and 1.4060, respectively. **b** Dependence of the calibration factor $\kappa_{\mathrm{EV}}$ for the contrast to size calibration on the presumed solid sphere equivalent refractive indices of the silica particles $n_{\mathrm{silica}}$ and EVs $n_{\mathrm{EV}}$.

### S2: Image pre-processing

The raw iSCAT frames $F'$ (200× average each) were flatfield corrected to reduce illumination inhomogeneities which otherwise overwhelmed the signal of captured EVs. Flatfield images $FF$ are median projections of 120 FOVs at random coordinates over the coverslip with 8× average each but otherwise identical imaging settings as $F'$. Each raw frame is flatfield corrected by pixel-wise division to yield normalized frames $F = F'/FF$. While the $F$ is void of large-scale illumination inhomogeneities (shown in Fig. 2**b** of the main text), variations on the order of a few percent often remained (e.g. inset in Fig 2**d**). These originated from variations in the LED intensity (single frame exposure of ~5 sec, flatfield generation takes ~180 sec) or from unique reflections at each position from the uneven plastic top of the flow cells which cannot not be fully removed by the flatfield processing. We therefore also applied a pseudo-flatfield[12] which is generated by applying a Gaussian blur with $\sigma = 10$ pixels on the frame $F$ itself to generate $F_{\mathrm{pseudo-FF}}$, and then update the original frame $F$ by $F$ / $F_{\mathrm{pseudo-FF}}$, analogous to the first flatfield operation.

Similarly, the fluorescence images were also corrected by a pseudo-flatfield. This was done since widefield images contained an overall fluorescence background from out of focus labels (for example from fluorescent EVs on the top of the flowcell) and because the fluorescence background could vary between first and second image (for example from bleaching the flow cell autofluorescence). We note that the pseudo-flatfielding is not as powerful as other lowpass image filters, however, it is fast and does not affect spot contrast values like, for example, image deconvolution. This step can be skipped if the images are taken in TIRF or confocal mode.

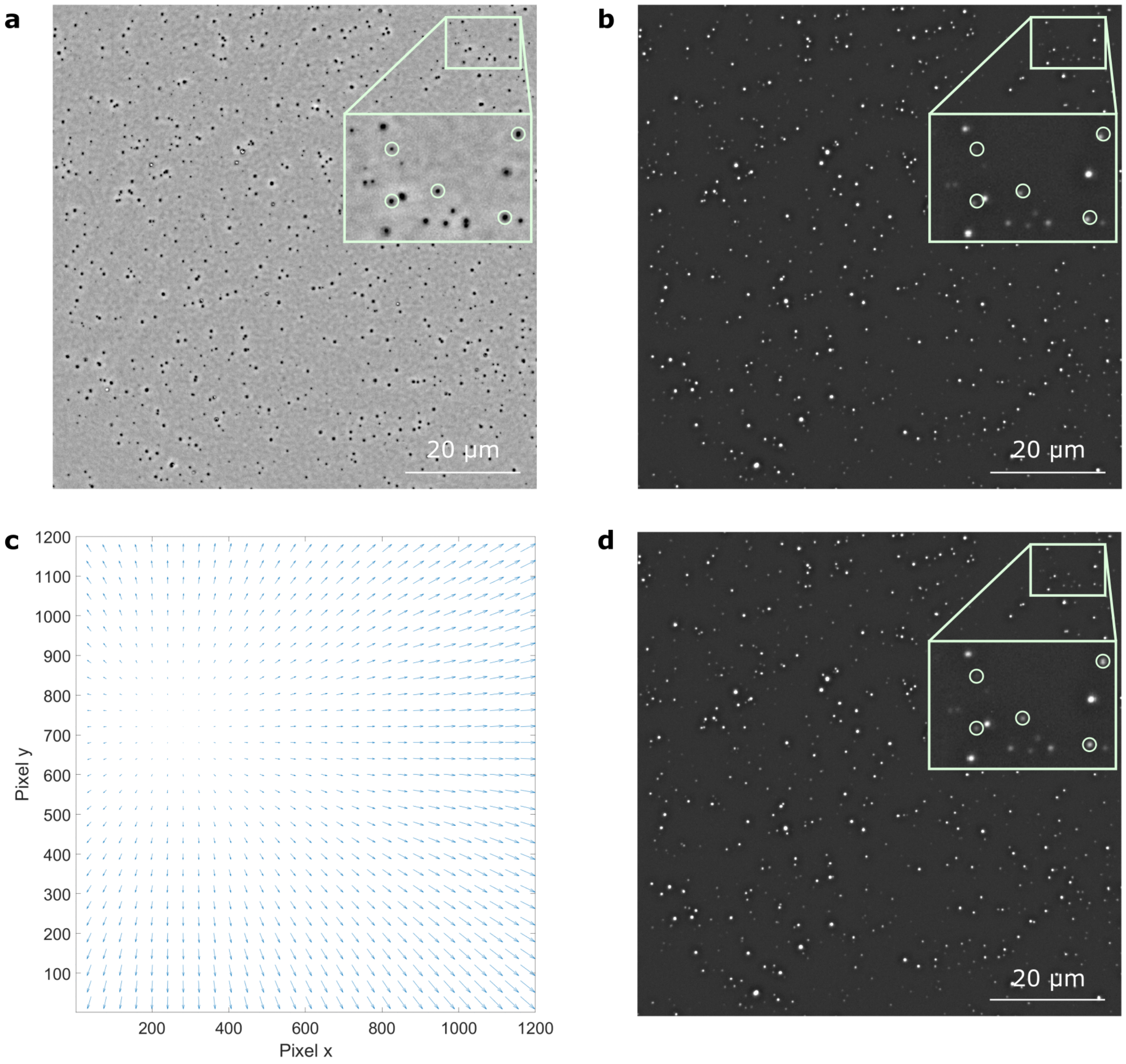

Figure S2: Chromatic aberration correction. Shown is a typical 1200×1200 pixel ROI (as used for all experiments in the main text) of an iSCAT image (**a**) and a fluorescence channel (**b**). The inset shows a matching region of interest in the top right image corner, where a few spots were marked based on their iSCAT signal. The inset in **b** shows the same spots and how they are slightly misaligned in the fluorescence channel due to chromatic aberrations. The actual algorithm selects only diffraction limited spots within a user defined contrast range. **c** Quiver plot mapping the chromatic aberration between the two channels shown in **a** and **b** over the 1200×1200 pixel ROI. The maximum displacement was ~5 pixel (ca. 370 nm), arrows are scaled for better visualization. **d** Processed fluorescence image where the inset shows the same ROI with spot markers as **a** and **b** and highlights the improved alignment in the final image illustrated by the central position of the fluorescent spot in the iSCAT circle iSCAT and fluorescent channels are furthermore affected by chromatic aberrations between each other which result in mismatches of several pixel between spot centers for overlaid images, shown in Fig. S3**a** and S3**b**. The maximum displacement vector between 440 nm iSCAT and the CTR channel in a 1200×1200 pixel ROI for example, is about 5 pixels, which is more than twice the diffraction limited spot size (for imaging with 100× objective with additional 1.5× zoom lens). These aberrations affect the accuracy of fluorescence measurements, for example only after the correction quadrant 2 on the top/right of the SPFI plots is identical whether the candidate spot localization is done of iSCAT or the fluorescence channel. Correcting aberrations between iSCAT channels is furthermore important when combining iSCAT contrasts from two channels into $C_{\mathrm{SP}}$.

One of the key function in our SPFI analysis pipeline is a pre-processing algorithm that can correct for chromatic aberrations between any two SPFI channels directly on the image data without use of special calibration samples. Since the distortion between various channels can have non-trivial shapes, we opted to map them on a pixel-by-pixel basis, Fig. S3**c**. Our algorithm made use of the fact that the EV image data contains many pairs of diffraction-limited spots which can be identified and matched automatically given that the capture density is low enough. The main idea is to collect the coordinates of matching pairs of spots over all FOVs, calculate the displacement vector (pixel-to-pixel mismatch) for all of them, and finally interpolate for all pixels of the full image. The first step is to localize spots in each channel individually and to select pairs of points with center-to-center distance of at most 10 while removing any spots with no match or cases in which multiple spots are found within 10 pixel distance. The resulting pairs of coordinates are then used to calculate an image displacement vector ($[u_{\mathrm{fluo}} - u_{\mathrm{iSCAT}}, v_{\mathrm{fluo}} -$

$v_{\mathrm{iSCAT}}$]) at the pixel coordinates $[u_{\mathrm{iSCAT}}, v_{\mathrm{iSCAT}}]$ of the reference spot in the iSCAT image. This was repeated in for all fields of view of the dataset (typically 20 - 30 fields of view). The resulting displacement vectors are combined and interpolated onto the full image grid where displacements for doubly sampled coordinates $[u_{\mathrm{iSCAT}}, v_{\mathrm{iSCAT}}]$ are averaged. Finally, the fluorescence image is transformed pixel by pixel to match the iSCAT as shown Fig. S3**d**. For further details on the algorithm, we refer the reader to the Matlab code (https://github.com/junckerlab/SPFI). Since the aberrations are setup specific yet identical for each measurement with a given channel combination, the correction map can be calculated once on a reference dataset and then be applied to all future datasets. We found it helpful to map the distortions of the full camera sensor and crop the map given the data ROI size.

### S3: Endpoint image alignment

To achieve efficient background removal for SP, it is crucial to ensure correct alignment between $F_{\mathrm{pre}}$ and $F_{\mathrm{post}}$. Small mismatches in the FOVs can readily arise from small holder movements during the pipetting during the particle incubation or imprecise stage movements between multiple FOVs and must be corrected digitally. The image alignment algorithm needs to be able to work on relatively few features that are present in $F_{\mathrm{pre}}$ (given that the glass substrate is usually clean) while tolerating the addition of many more particles in $F_{\mathrm{post}}$ from the sample incubation. The first step of the algorithm is to localize diffraction-limited spots within a manually set contrast range in both iSCAT incubation images by a simple local minima search. We denote the coordinates in the pre-image $\overrightarrow{u_k}$ and the localized spots in the post-image $\overrightarrow{u_l}$. Then, the displacement vector $\overrightarrow{d_{kl}}$ of each spot $k$ in $F_{\text{pre}}$ to all spots $l$ in $F_{\text{post}}$ is calculated, $\overrightarrow{d_{kl}} = \overrightarrow{u_k} - \overrightarrow{v_l}$. All matching spots in the two images produce the same displacement vector $\overrightarrow{d_{kl}}$, regardless of their actual positions, whereas non-matching spots produce random $\overrightarrow{d_{kl}}$. The correct image mismatch $\vec{d}$ is thus simply found by searching for the most common $\overrightarrow{d_{kl}}$ . The algorithm generates a two-dimensional histogram of all displacement vectors with 4 by 4 binning (to account for minor pixel shifts in spot centers), searches for the maximum bin to find $\vec{d}$, and finds the correct displacement from the average displacement in the winning bin. Further details can be found in the Matlab code (https://github.com/junckerlab/SPFI).

We note that if too few features are available in $F_{\text{pre}}$, e.g. on completely clean glass slides, a small amount of silica or gold particles can be added to act as reference. As little as 5 particles in $F_{\text{pre}}$ were found to be sufficient among >>100 of additional particles in $F_{\text{post}}$ for the algorithm to work in a fully automated and robust way.

### S4: Candidate spot localization:

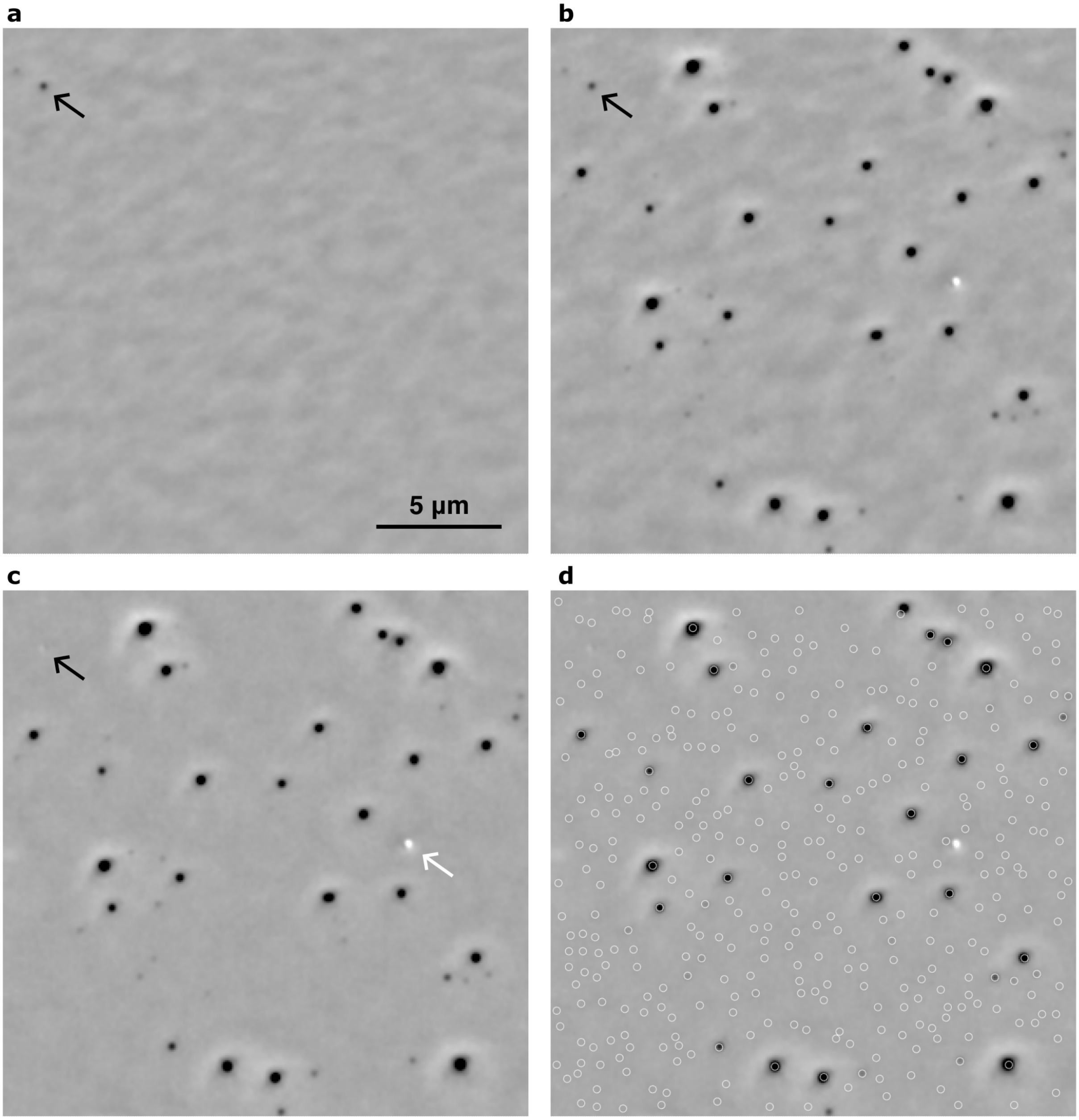

Figure S4: Selection of candidate spots in the iSCAT difference images demonstrated on EVs from HT29 cells. Shown are $F_{\text{pre}}$ (**a**), $F_{\text{post}}$ (**b**), $F_{\text{diff}}$ (**c**) and $F_{\text{diff}}$ with all candidate spots marked as circles (**d**). The black arrow shows a high contrast spot which is not fully removed in the difference image **c** but can readily be identified in in the pre-incubation image **a** alone and then be masked for the candidate localization. The white arrow indicates a bright spot (large EV) which is not registered by the contrast minimum search as candidate spot but could potentially be included with dual wavelength iSCAT imaging with a longer wavelength.

The first step of our particle registration algorithm is the registration of background impurities that are present before particle incubation to establish a reproducible noise floor for the SPFI plots. Taking the difference between pre- and post-incubation images $F_{\mathrm{diff}} = F_{\mathrm{post}} - F_{\mathrm{pre}}$ was efficient at removing small particles and glass roughness. Larger dust, however, often had residual contrast in $F_{\mathrm{diff}}$ that was comparable to the smallest captured particles (see black arrow in Fig. S4). This was mainly caused by small microscope focus drifts between $F_{\mathrm{pre}}$ and $F_{\mathrm{post}}$ are imperfect image alignment (even when shifting with subpixel accuracy). To remove those spots without purely relying on image subtraction we applied a simple crud registration directly on $F_{\mathrm{pre}}$ and masked the affected areas for the candidate localization.

The spot localization algorithm then registers so-called candidate spots on $F_{\mathrm{diff}}$ by simply registering all local contrast minima (i.e. all pixels which are darker than all their eight neighboring pixels) as shown by the white circles in Fig. S4**d**. This algorithm does not include any bright spots (see white arrow in Fig. S4**c**), as they cannot be sized based on our $d^3$ model. Whenever iSCAT imaging is done using 440 nm and 740 nm illumination, candidate selection is first done on the 740 nm channel and only spots exceeding a contrast threshold, typically $\sqrt[3]{C_{740\,\mathrm{nm}}} > 0.2$, are kept. A second round of candidate localization is then done on the 440 nm channel, and both sets of candidate spots are combined to ensure both high and low contrast spots are included.

The last step of the candidate localization algorithm is to measure iSCAT contrasts and florescence intensities in all recorded channels as single pixel intensity measurements. Before doing so, each channel is blurred using a Gaussian filter with sigma of one pixel to remove uncorrelated pixel noise. The kernel size corresponds to about 2/3 of the microscope PSF size (at 440 nm using the 100× objective with 1.5× additional zoom lens) and is thus barely affecting peak contrast values. Once again, further details can be found in the Matlab code (https://github.com/junckerlab/SPFI).

We note that two variants of the candidate spot localization are possible, either the iSCAT is used to register the EVs and then the fluorescence signal is recorded subsequently, or vice versa. The algorithm for each case is analogous and simply differs in whether candidates are registered as local contrast minima or fluorescence intensity maxima. While top/left or bottom right quadrants of the resulting scatterplots might differ due to stochastic sampling of the background, in our experience the top/right quadrant was conserved between the methods given that the images were corrected for chromatic aberrations.

Candidate spots include the EVs but deliberately also include low contrast features from the image background. The addition of these background spots means the scatterplot data is only minimally processed and different datasets can be readily compared since variations in background due to low quality substrates or non-specific binding of fluorescence labels are not excluded from the plots but simply fall into the top/left or bottom/right quadrants of the scatterplots in buffer controls. Furthermore, downstream data processing can leverage established flow cytometry workflows which also include noise and signal data and use combinations of label-free and fluorescence signals to gate the particles of interest.

The candidate registration algorithm based only on local minima (or maxima for fluorescence images) of the image data was chosen both for its simplicity and robustness. On the downside, it does not consider that the EVs are mostly diffraction limited spots and as such includes an excessive number of spots at the image noise floor (see also Supplementary Section S6). We performed tests where we thresholded candidate spots based on their spot circularity, e.g. by difference of Gaussian or radial variance transform algorithms. We found that additional thresholding on circularity did significantly reduce the number of candidate spots in the image noise (quadrant 3 in the bottom/left of SPFI plots) and was able to remove some candidates localized in minor image artifacts (c.f. Fig. 2d of the main text). However, it required at least one additional user-defined threshold value and furthermore did not change any findings from the SPFI plots of the main text in a meaningful way. While we plan to include additional thresholding in future studies, we did not use it for any data shown in this manuscript.

### S5: Dynamics of particle capture

Conveniently, SPFI does not require precise knowledge on the concentration of EV samples since the EV immobilization on the capture surface can be monitored in real-time as shown in Fig. S5**a**. We show real-time recordings for HT29 EV capture on PLL surfaces in Supplementary Video 1 and capture on glass in Supplementary Video 2. Both videos were flatfield processed and all frames are 4× averaged. The incubation can be stopped once a desired density is reached while still ensuring sufficient separation for recognition between stochastically distributed EVs. Since most small EVs are diffraction limited spots in iSCAT, they can, in principle, be distinguished down to the Abbe diffraction limit $d_{\min} = \frac{\lambda}{2\,NA} \approx 150$ nm ($\lambda = 440$ nm, $NA = 1.45$) where they produce two distinct contrast minima in the candidate search. However, spots in the iSCAT image also must be matched to fluorescence images and, despite image pre-processing, there can be residual chromatic aberrations between the channels. Furthermore, there are situations in which fluorescence spots do not coincide with the EV centers as some protein labels (e.g. CD63-GFP) might be clustered on one side of the EV. Upon inspection of the datasets, we estimated that the offsets between iSCAT and fluorescence channels fall within at most three times the resolution limit, or $3d_{\min} \approx 450$ nm. S5**b** shows numerical calculations to estimate the maximal capture density for both the above limits using a Monte Carlo method. We simulated a capture area of $(100\ \mu\text{m})^2$ and added particles one by one at randomly generated coordinates. In each step, the distance between all the particles was calculated and the total number of particles without any partner within $1d_{\min}$, $2d_{\min}$ and $3d_{\min}$ were counted. The simulation was performed 100 times, and the results were averaged in Fig. S5**b.** To establish a maximum capture density, we required 95 % of spots to be isolated within $3d_{\min}$ such that their contrast and fluorescence signals are guaranteed to be matched correctly (dashed gray line). We found that this threshold was crossed at 792 particles per (100 μm)$^2$ or equivalently at 614 particles per (88 μm)$^2$ which is the FOV size used for all SPFI data of the main text. Imaging in 20 FOVs per flow cell thus allowed us to analyze ~11,000 single EVs as reported in the main text.

It must be noted that the 95 % threshold does not necessarily imply a 5 % error rate for SPFI since likely only a fraction of these spots will be mismatched between iSCAT and fluorescence channels. Furthermore, with 792 EVs per (100 μm)$^2$, only about 0.6 % of vesicles had a partner less than $1d_{min}$ apart and are therefore considered truly indistinguishable. Most spots within the $3d_{\text{min}}$ limits are still distinguishable in iSCAT alone and can in principle be removed from the candidates, for example by simply considering spot circularity as mentioned in section S4.

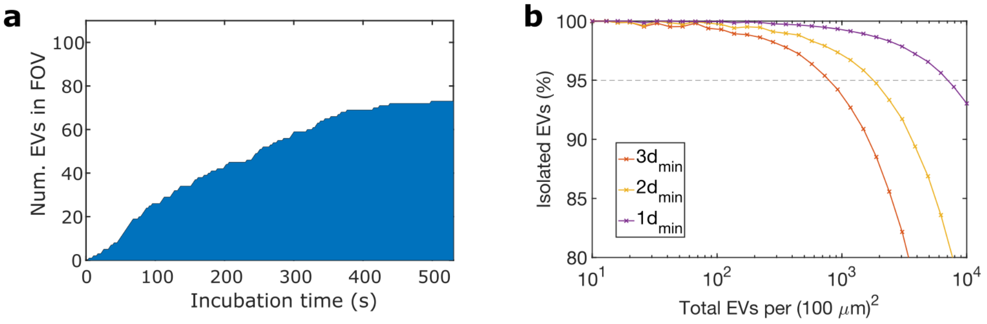

Figure S5: Particle capture and optimal packing density. **a** Time trace of the cumulated number of particles captured during incubation. At time zero, purified EVs from HT29 cells at approximate concentration $10^9$parts/ml were flushed into the flowcell. A timeseries experiment was started in which a region of interest of 512 by 512 pixels (ca 38 × 38 μm$^2$) was recorded at 100 FPS. During the analysis, 200 frames were averaged and the EVs that were deposited were detected using the algorithm described above. **b** Monte Carlo type numeric calculation to estimate the maximum packing density for EVs in a 100 × 100 μm$^2$ area. We assigned random EV positions and calculated the number of isolated EVs without any partner within multiples of the Abbe resolution limit $nd_{\text{min}} = 150$ nm where $n = 1,2,3$. Requiring that 95 % of EVs have no partner within $3d_{\text{min}}$, results in a maximum density of 792 particles per (100 μm)$^2$.

### S6: Size photometry noise floor

While the noise floor in iSCAT is given by the shot noise and technical readout noise (Fig. 2**c** of the main text), it was found that real SP experiments produced a slightly enlarged 'noise blob' (Fig. 2**d** of the main text). The principal reason is that in our noise estimate (Fig. 2**c**) we calculated the standard deviation of pixel-to-pixel image noise whereas SP data shows candidate spots registered as local contrast minima and the single pixel contrast measurements thus sample the extremes of the noise distribution. Refined candidate spot localization or contrast measurement techniques (e.g. Gaussian fits) could alleviate this excess. Additionally, however, pre- and post-incubation images often contained several characteristic artifacts shown as inset in Fig. 2**d** of the main text that further increased SP noise. These were identified as out of focus scattering from dust from within the microscope body which was not efficiently removed by flatfield processing. The main reason for this was that the dust particles seemed to be in the illumination arm of the microscope such that they scatter light which gets reflected from the glass coverslip before reaching the camera. If the coverslip surface is slightly bent, either concave or convex, the scattered light is reflected slightly off-axis. This means that the image of the dust particle is not static on the camera sensor when the flatfield image is taken at multiple positions with different coverslip curvature. We tested SP on two independent microscopes and found similar image artifacts on either one. None of these artifacts were apparent in other imaging modalities such as fluorescence or brightfield imaging as there is no reflective glass surface. It was found that the severity of the artifacts could be limited by taking great care that glass substrates (type 1.5 cover glasses which are somewhat flexible) were mounted as flat as possible. Since these features appear roughly in the same positions (beyond the few pixel shifts), they can in principle be cut out automatically and not considered for further analysis.

### S7: iSCAT particle contrast simulations:

We performed several FEM simulations for silica particles and EVs to further investigate the iSCAT contrast switch and estimate the SP sizing limit. Fig. S6**a** shows the simulation geometry implemented in the COMSOL *electromagnetic waves, frequency domain (ewfd)* module consisting of bottom and top regions modeling the flat glass substrate ($n_{\text{glass}} = 1.51$) surrounded by water ($n_{\text{water}} = 1.33$). A planar electromagnetic wave modeling the illumination is injected through input port 1 on the bottom. It is partially reflected by the glass substrate and otherwise transmitted to output port 2 on the top. Periodic Floquet conditions were applied in both *x* and *y* directions to reduce computational load. We first verified that the reflected power in port 1 without any particle was ~0.36 %, as expected by the Fresnel law. Next, we modeled the silica particles as solid spheres of radius $r$. The sphere centers were placed at a $z$ height of $r - 5$ nm and the bottom of the spheres below $z = 0$ was removed. This was done to avoid the sphere touching in only a single point which would lead to simulations not converging and does only marginally affect the results. The electromagnetic fields were then solved and the total reflected power in port 1 was calculated. The result was compared to a second simulation where the particle refractive index was set to water (to avoid changes in geometry or meshing) such that a reference value for the reflected power from the substrate could be obtained, see Fig. S6**b**. The iSCAT contrast was calculated by dividing the reflected power of both simulations to estimate the on-axis contrast. We note that the FCM simulations go beyond the standard analytical theory of section S1, however, they remain an approximation of the actual measurement process. In particular, they still do not consider many aspects of the microscopic detection such as objective collection angles, point spread function, aberrations[13] or phase factors due to the Gouy phase[8].

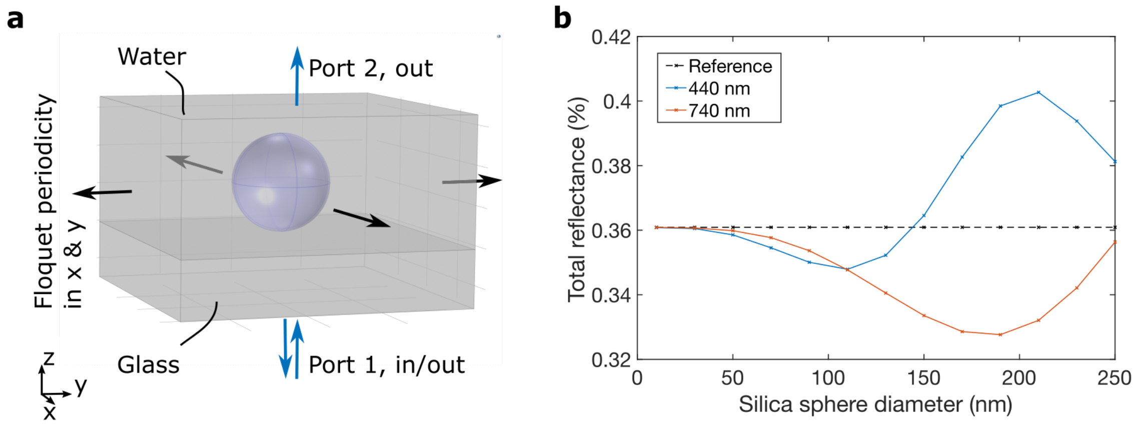

Figure S6: **a** Geometry of COMSOL models to simulate iSCAT contrasts of silica particles and EVs. **b** Simulation results for silica particles under 440 nm and 740 nm iSCAT illumination. Simulations were performed twice for each particle size, once with the refractive index of the sphere set to the particle of interest and once to the refractive index of the surrounding medium. The total reflected power in port 1 of the simulation geometry was measured each time and the two values were divided, mimicking the experimental definition of SP contrasts which are relative to the background. Note that COMSOL correctly predicts a reduction of total reflectance due to the interferometric effect for small particles (i.e. iSCAT produces dark spots), but that the SP contrasts throughout this manuscript are defined as $C = 1 - \frac{P_{\mathrm{int}}}{P_{\mathrm{ref}}}$ giving positive values.

The first set of COMSOL simulations was performed for silica particles, where we used refractive indices of $n_{\mathrm{Silica}} = 1.4663$ at 440 nm and $n_{\mathrm{Silica}} = 1.4544$ at 740 nm illumination wavelength[9]. The width of the simulation geometry in $x$ and $y$ was set to 1 µm, the thickness of simulated glass below the particles was 350 nm and the height of the water domain was 500 nm. We verified that increasing the domain sizes did not change the shape of the simulated contrast curves. However, increasing lateral widths reduced the interferometric contrast relative to background since they were calculated from the total reflected power and increasing the simulation geometry means the glass-water interface contributes more relative to the particle. This furthermore means that the simulation results are not absolute and had to be scaled to match the experimental contrasts in magnitude. We thus applied setup calibration factors $\nu_\lambda$ to the simulation results and used least-square fits to the measurements for 50 nm and 80 nm particles for 440 nm and to measurements for the 50 nm, 80 nm and 100 nm particles for 740 nm wavelength to determine $\nu_{440\,\mathrm{nm}} = 1.32$ and $\nu_{740\,\mathrm{nm}} = 0.82$. These factors were applied to the simulation data of Fig. 4 of the main text and furthermore to Fig. S7 below to ensure simulations and experimental data can be compared directly.

As discussed in the main text, we found that experimentally measured liposome and EV contrasts at 440 nm illumination could exceed 5 % whereas COMSOL simulations with spherical EVs predicted maximal values well below 2 % before the contrast inversion. We therefore performed further COMSOL simulations to investigate if a partial 'collapse' of soft particles upon adsorption to the glass substrate could explain this observation**.** To model the flattening of spherical particles with diameter $d$, we generated ellipsoids ($n_{\mathrm{ellipsoid}} = 1.37$) whose vertical axes were squeezed to 100%, 66% and 33% of $d$ while expanding in the horizontal semi-axes to maintain the volume of the respective spherical particle (which may not reflect reality as water and small molecules may escape from squeezed vesicles). The resulting contrasts at 440 nm are shown in Fig. 4**b** of the main text. We found that all curves overlap for small vesicles below about 100 nm, and that the contrast inversion is delayed further the more the ellipsoids were squeezed. For ellipsoids with 2/3 of the original height, we found that maximum contrasts were above 4 %, and that the inversion point for the ellipsoids with 1/3 height was delayed to outside our simulation range with maximally simulated contrasts exceeding 20 %. We then performed a least-square fit of the data for the 1/3 height ellipsoids below 200 nm to the $\propto d^3$ model and found that the simulation data closely followed the $d^3$ curve. Squeezing of liposomes or EVs upon capture could thus explain the experimentally measured iSCAT contrasts since it extends the range in which SP using the $d^3$ law holds true. Importantly, it did not seem to affect the values at smaller $d$, which still gave the same contrast values as the spherical particles.

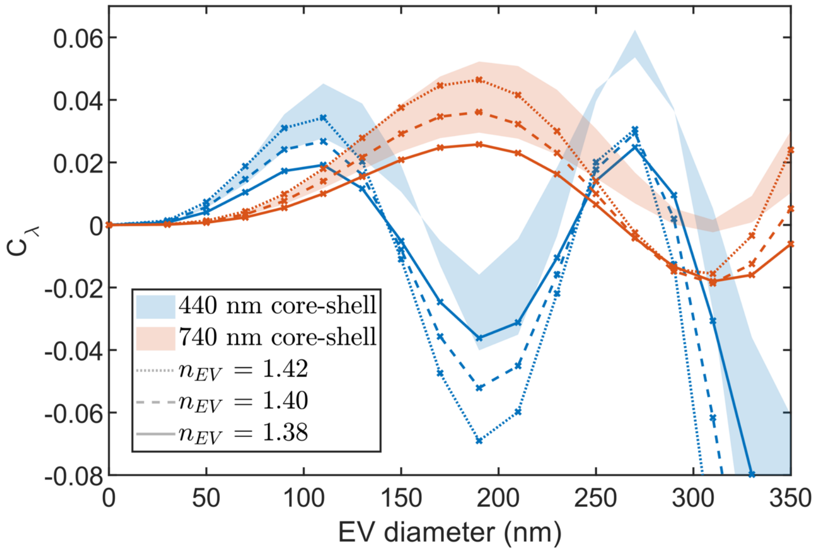

Figure S7: COMSOL simulations comparing SP contrasts of solid sphere EV models and core-shell EV models to complement Fig. S2**a**. Solid sphere equivalent refractive indices of $n_{\mathrm{EV}} = 1.38$ (solid lines), $n_{\mathrm{EV}} = 1.40$ (dashed lines) and $n_{\mathrm{EV}} = 1.42$ (dotted lines) are shown whereas the core-shell geometry was simulated twice for different optical densities of the EV core (see text) and the region between the two results was shaded. All curves reproduced the initial $d^3$ scaling (verified by least-square fits, not shown) and only differed in magnitude. At larger vesicle diameters, the models showed larger discrepancies of both magnitude and location of the contrast inversion.

The $d^3$ law for sizing in SP was validated on solid silica spheres with homogenous refractive index, however, EVs have optically denser membranes compared to their lumen. This results in a size dependence on the overall solid sphere equivalent refractive index of EVs which should be considered for sizing EVs over a large range of diameters. We performed additional simulations to compare solid EVs with core-shell EV models to validate that both models reproduce $d^3$ contrast behaviors for small particles. The core-shell geometry was implemented in COMSOL using two spheres of radii $d/2$ and $d/2 - t$ to model the EV membrane (refractive index $n_{\mathrm{membrane}}$) of thickness $t$ surrounding the vesicle lumen (refractive index $n_{\mathrm{lumen}}$). We used the default values implemented in FCMPass[14] v. 4.1.1 such that the SP calibration is comparable to the FCM calibration (Fig. S14). The shell thickness was set to $t = 5$ nm and the shell refractive indices to $n_{\mathrm{shell}} = 1.4863$. The low refractive index models used a core refractive index of $n_{\mathrm{core}} = 1.3658$, whereas the high refractive index models used $n_{\mathrm{core}} = 1.4060$.

The comparison is shown in Fig. S7 where the range of the core shell model is indicated as shaded areas for both wavelengths, which is contrasted with three homogeneous models with solid sphere equivalent refractive indices of $n_{\mathrm{EV}} = 1.38$, 1.40 and 1.42. We found that all models reproduced the $d^3$ contrast scaling for low EV diameters, which was verified by least-square fits to initial datapoints (not shown). Beyond the contrast inversion points, the simulated contrasts between homogenous and core-shell models differed substantially. We found that the solid sphere equivalent refractive index of $n_{\mathrm{EV}} = 1.40$ best approximated the core-shell models in 440 nm and 740 nm imaging simultaneously in accordance to Fig. S2**a**.

### S8: Particle contrasts with small defocus

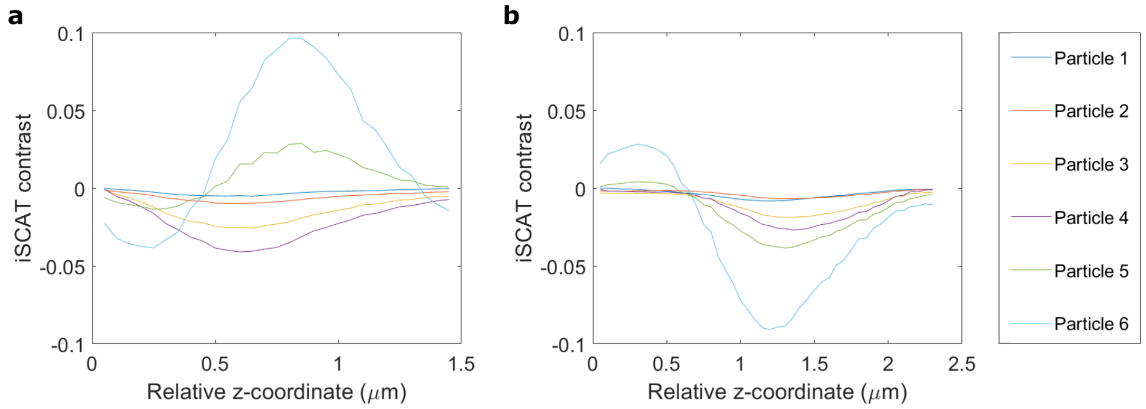

Figure S8: Contrast vs. defocus for several silica particles in a mixed population of 20 nm, 50 nm, 80 nm, 100 nm and 200 nm particles imaged in the same flow cell (note that aggregates of the smaller particles, e.g. 20 nm, are indistinguishable from single larger ones). iSCAT contrast of particles as function of *z*-coordinate with arbitrary offset relative to the surface with **a** 440 nm illumination and **b** 740 nm illumination. The iSCAT contrast values are relative contrast to the background surrounding the particles, negative values here imply ‘dark’ iSCAT contrast. We found that as the minima of the curves are getting lower, they are also tended to shift to the left. This is caused by an interplay of Gouy phase and the fact that optimal focus plane, presumable the center plane of the particles, is shifted for particles of varying diameters.

### S9: Additional SPFI experiments on EVs

This section contains several control experiments on EV samples that were performed in support of the data shown in Fig. 5 of the main text.

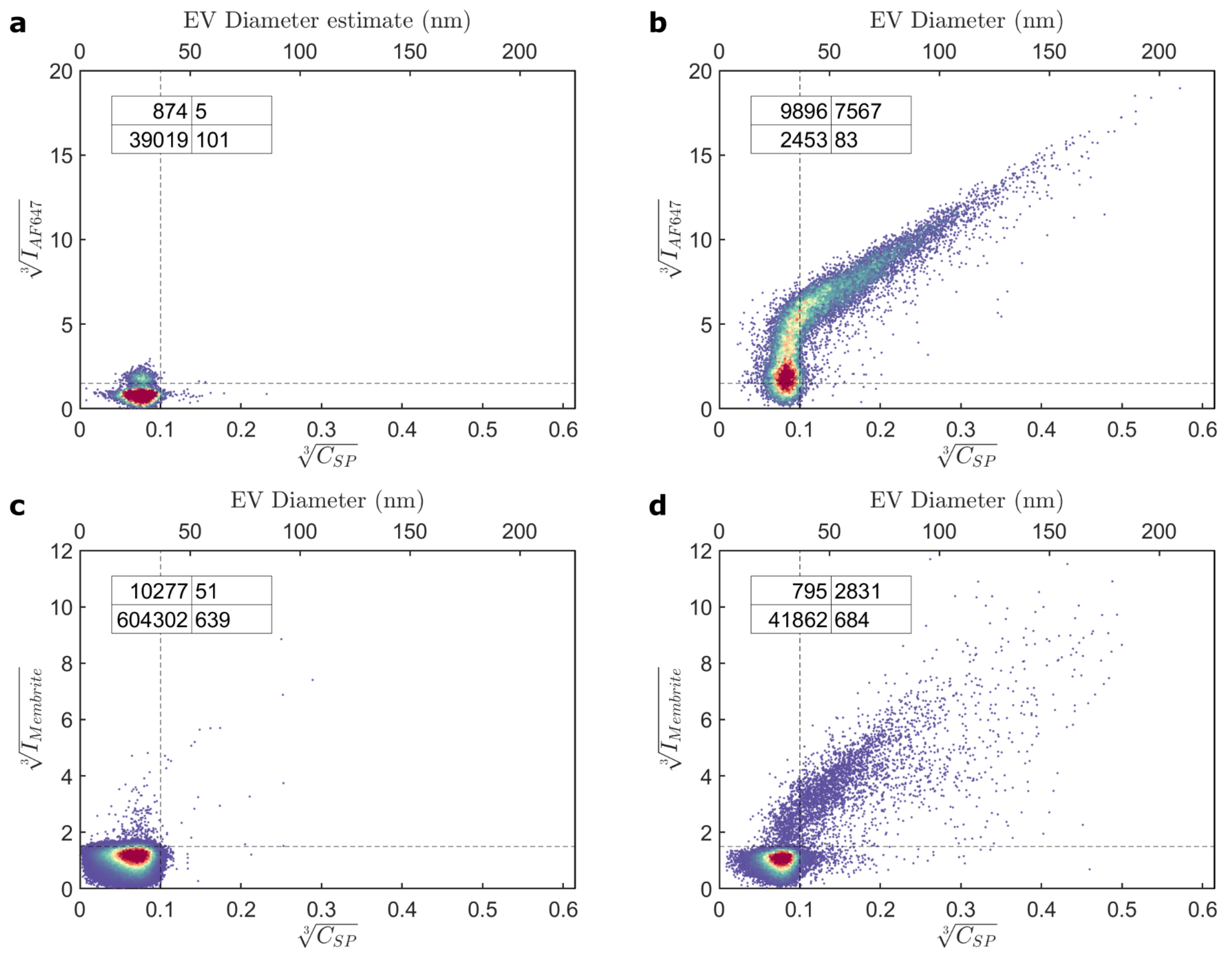

Fig S9: Additional data on membrane staining to support Fig. 5**a**. **a** PBS biotinilation control and **b** HT29 derived EVs which were biotinylated, stained with AF647 conjugated streptavidin, and captured on PLL coated glass surfaces. The points in quadrant 1 on the top/left of **a** is signal from individual AF647 conjugated streptavidin which, due to its relatively high molecular mass of ~60 kDa, was not completely removed during purification using the 100 kDa Amicon filters (see methods). **b** Biotinylated HT29 EVs, same data as Fig. 5**a** of the main text. **c** MemBrite488 staining control and **d** MemBrite488 stained HT29 derived EVs. MemBrite staining is noisier than biotinylation, but the sub-linear scaling for the maximal fluorescence intensity (when plotted as third root intensity) as functions of size persists.

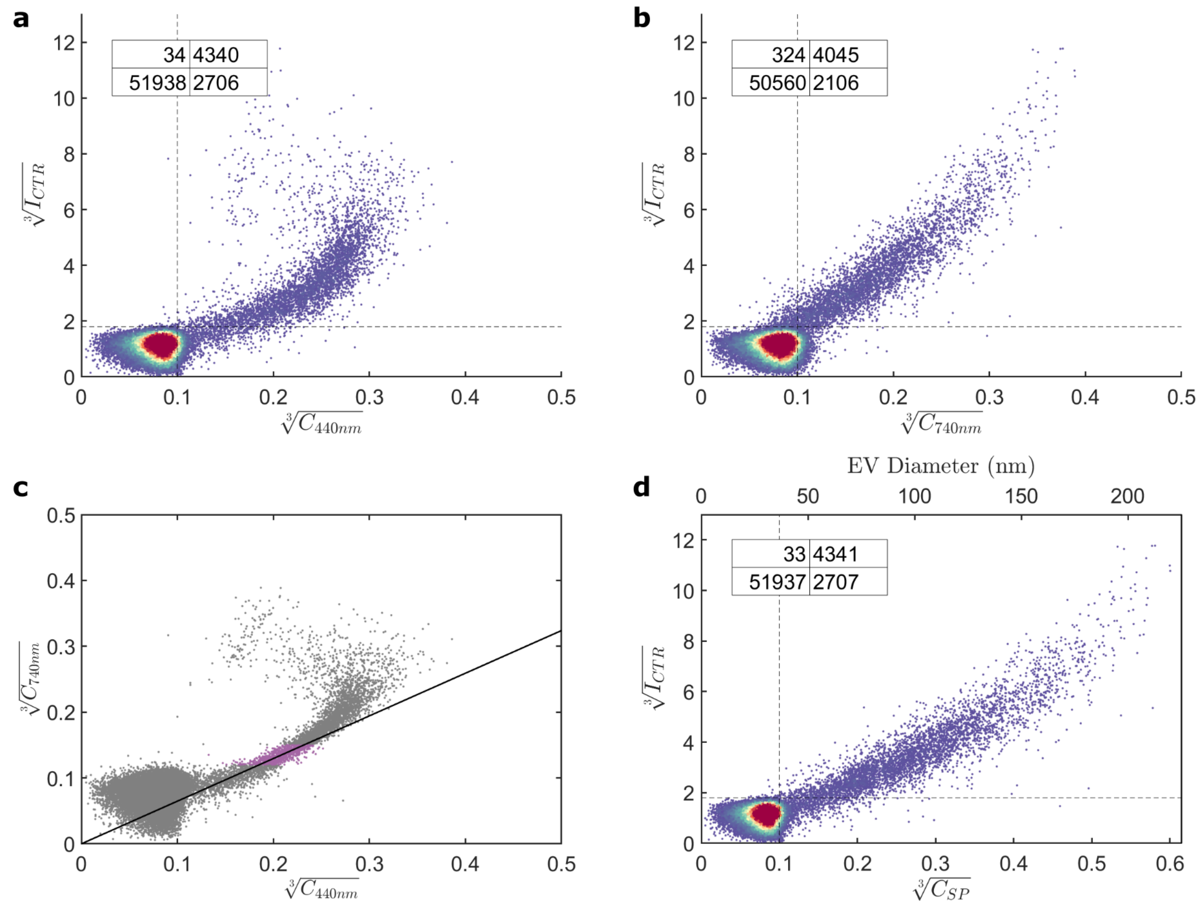

Fig S10: Individual SPFI plots of the CTR stained EVs shown in Fig. 5**b** of the main text and the combination into a combined SP contrast. **a** Imaged in 440 nm illumination. **b** Imaged in 740 nm illumination. EVs in the sample clearly exceed the SP range in 440 nm (**a**) as seen by the bend in the distribution whereas the 740 nm (**b**) contrasts are still approximately proportional to the CTR fluorescence intensity. **c** SP correlation plot which allows to fit linear part of the distribution (the black line is a least square fit to the purple points) such that the two sizing ranges can be combined. **d** Combined SP contrast measurement as shown in Fig. 5**b** of the main text.

Fig. S10 shows an independent experiment in which HT29 EVs were co-stained with CTR and Carboxyfluorescein succinimidyl ester (CFSE) to compare these two EV stains. The EVs in this experiment were purified identically to the other samples (see methods) except that the supernatant was filtered with 0.22 µm filters instead of 0.45 µm filters and iSCAT imaging in 440 nm was found to be sufficient for SP. Staining for Fig. S10 was done by incubating 0.5 ml of SEC purified EVs (concentration ~$10^9$ particles per ml measured by NTA) with 0.5 µl of 1 mM CTR and CFSE solution in DMSO for two hours at 37 °C. The unbound stain was washed six times (resulting in a dilution of $10^6$) using a 0.5 ml 100 kDa Amicon filter (each wash consists of adding 500 µl PBS, spinning 5 min at 5,000 g and discarding the flowthrough volume). Staining was performed twice for all samples and controls to maximize the fluorescence signal. The incubation times for all samples were matched to 3 min.

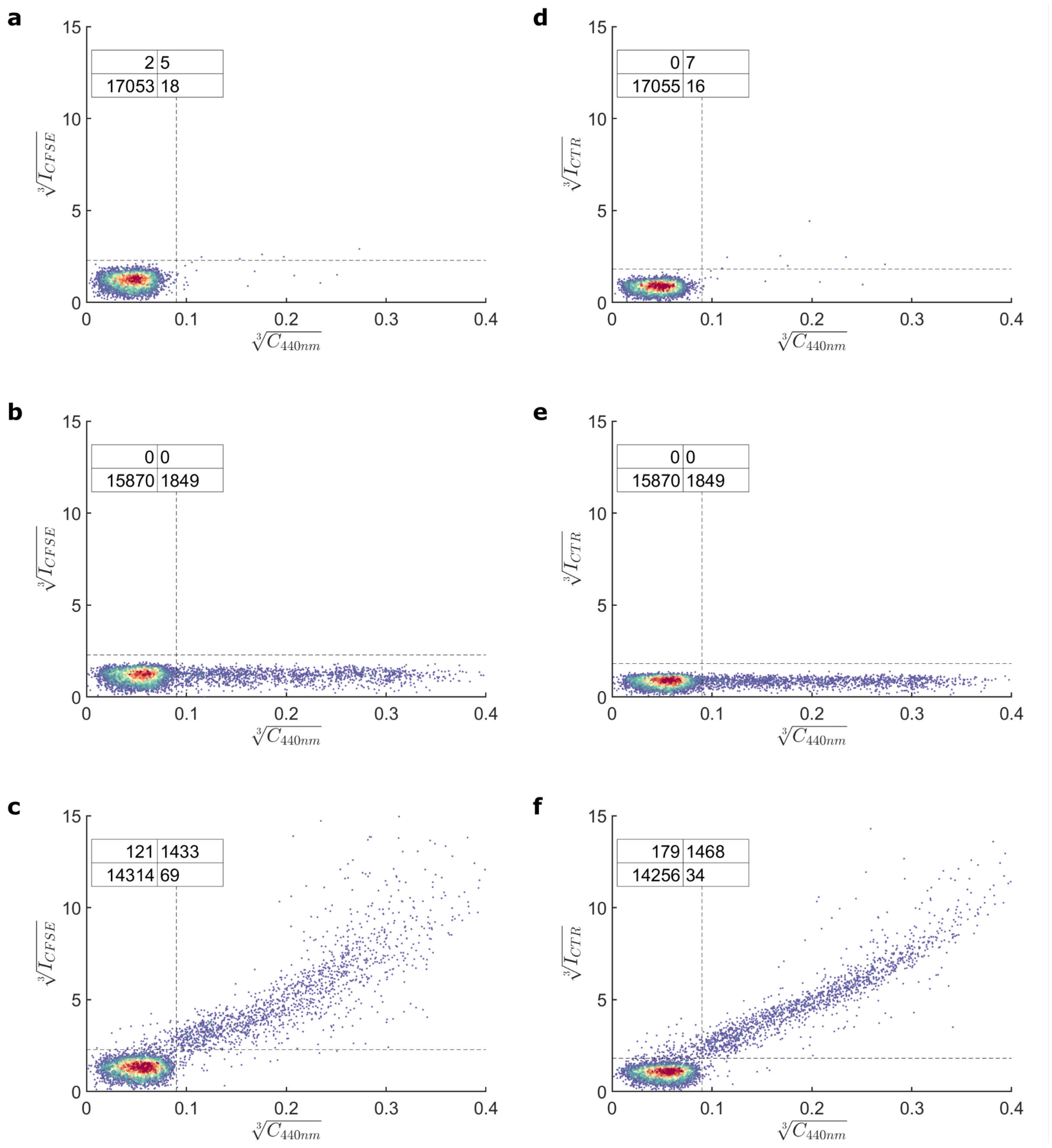

Fig S11: Additional experimental data on CFSE and CTR stained EVs comparing negative controls of ‘stained’ buffer and unstained EVs to actual stained EVs. **a** CFSE stained PBS, **b** unstained EVs, **c** CFSE stained EVs, **d** CTR stained PBS, **e** unstained EVs, **f** CTR stained EVs. All data was generated by registering candidate spots in the 440 nm iSCAT channel and measuring the fluorescence intensities subsequently. Insets indicate the number of spots from the scatterplots that fall in the respective quadrants of the gates (dashed lines).

Fig. S12 shows a comparison of different measurements performed on the same sample of A431 derived CD63-GFP EVs which is also used in Fig. 5**c** of the main text. SPFI data was taken once after capture on positively charged PLL (Fig. S12**a**, 12**c**) and once after capture on plasma activated, negatively charged bare glass (Fig. S12**b,** 12**d**). Given the faster capture on PLL, the sample was diluted 1:5 on PLL whereas the stock concentration was used on glass, and both incubation times were approximately 30 s. We found that the SPFI plots show minor differences but that the two clusters of points described in the main text are readily distinguishable. A full investigation on the effects of surface choice regarding possible selection bias or effects on SP sizing will be part of future work. Figs. S12**e**, S12f show FCM buffer control and sample measurements on the same EVs where the GFP signal is recorded as height (H) of the signal in the FITC channel. Fig. S12**g** shows the size distribution of the sample measured by NTA via scattering detection.

Fig. S12**h** combines the recorded size distributions from SP (on glass), FCM, and NTA measurements in a single plot for visual comparison of the respective sizing ranges. The data is plotted in log scale for visibility and each histogram is individually normalized by the height of the histogram bin centered at 130 nm which allows to overlay the curves and compare the acquired distributions. Among the three methods, SPFI has the lowest LOD (note that the highest histogram bars correspond to the background noise seen in the bottom left of Fig. S12**a**, S12**b**) but also the lowest upper sizing limit which is given by the contrast inversion in the 740 nm imaging channel. We note again that the contrasts are calibrated from the silica size standards under the approximation that all EVs have the same refractive index of $n_{\mathrm{EV}} = 1.40$, neglecting size dependence (Fig. S2**a**) and random refractive index variations from protein density.

FCM sizes EVs in solution by side scattering detection with a lower detection limit around 100 nm after calibration, as illustrated in Fig. S14. This calibration mirrors the one for SPFI, where various polystyrene beads of known size and refractive index are measured to fit the detector response function in FCMpass[14], thereby allowing to estimate EV sizes by approximating their refractive indices. Given that FCM has a larger sizing range compared to SPFI (and that the technique is much more mature) the refractive index size dependence is explicitly considered in the calibration. FCMpass offers three predictive models corresponding to low, medium, and high refractive index scenarios and we used the one for medium refractive index. Variations in EV refractive index caused by protein density, however, cannot be accounted for during the calibration, which leaves a similar caveat in FCM as in SP sizing. It is also worth mentioning that FCM analysis can encounter plateaus in detector response. For our system this occurs around the 400 nm – 500 nm size range as shown in Fig. S14, where sizing becomes less accurate (see also Fig. S13). These plateaus result from Mie resonances traversing the detector acceptance angles and are specific to each instrument.

NTA nominally has the largest detection range from ~50 nm to about 1 $\mu$m specified by the manufacturer. The practical dynamic range for sizing polydisperse samples such as EV, however, is substantially smaller. Firstly, the lower detection threshold is characterized by a gradual decrease in sensitivity from particles sized 100 nm down to approximately 50 nm. This is because smaller particles diffuse faster than larger ones and might leave the detection zone faster than the minimum tracking length can be acquired. This means small particles in polydisperse samples are underrepresented and NTA curves produce a characteristic peak at intermediate EV sizes (here at ca 90 nm) instead of following the inverse power law size distribution to much smaller EVs[15]. The limit on dynamic range for polydisperse samples occurs because camera overexposure from the strong scattering of large particles needs to be avoided (max. 20 % of overexposure according to the manufacturer) which, at maximum sensitivity to small EVs, limits the upper sizing range in our measurements to around 300 – 400 nm. Indeed, larger EVs (at low concentration) can be found when reducing the camera level. We note that NTA measures hydrodynamic EV diameters in contrast to SP and FCM which both measure physical EV diameters. Hydrodynamic diameters depend on the physical EV size but can extend significantly due to hydration shells that form around EVs and other particles from the interaction with the immediate environment. A notable advantage to NTA is that hydrodynamic diameters can be calculated from particle tracking data without calibration and without relying on reference samples.

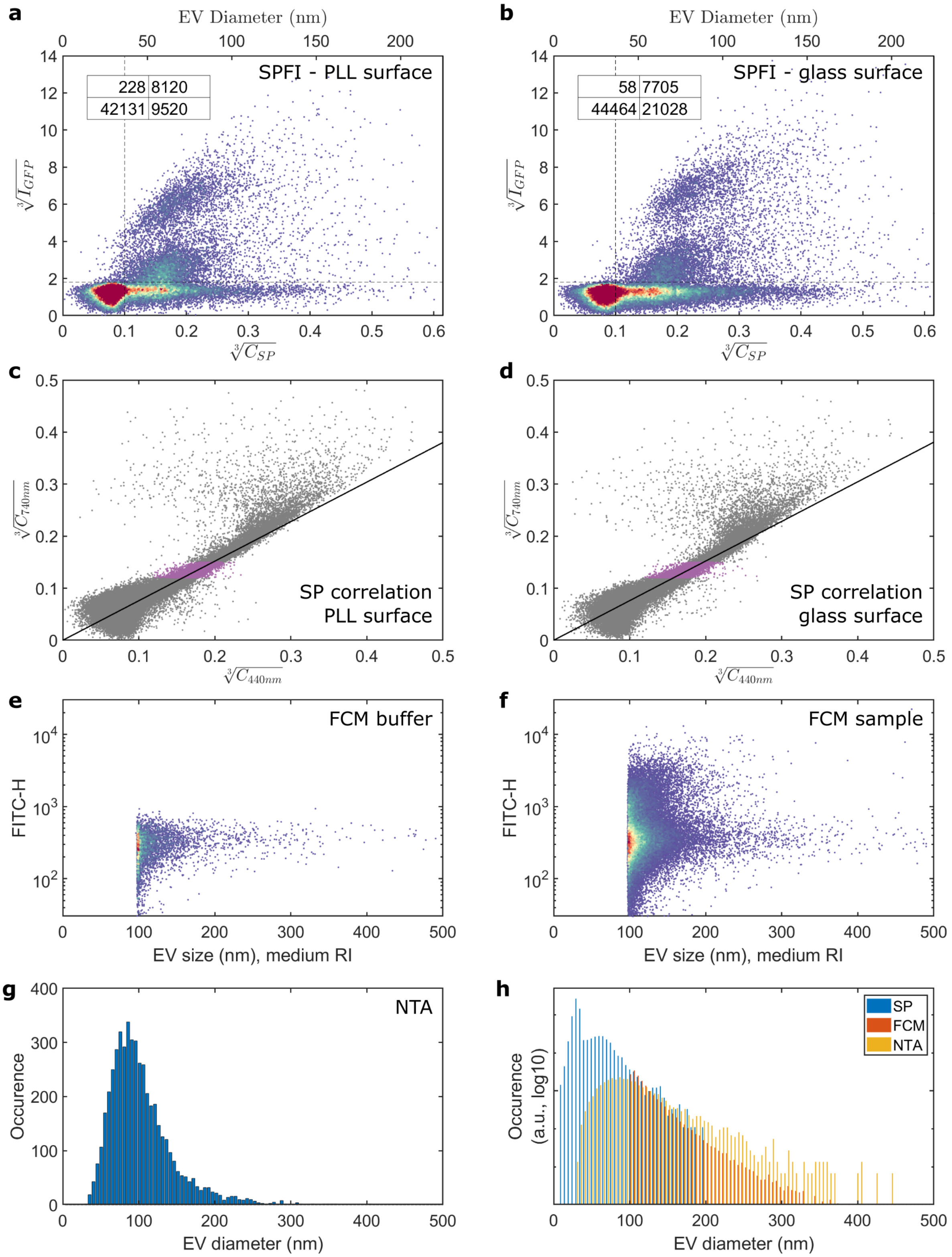

a
EV Diameter (nm)
SPFI - PLL surface
228 8120
42131 9520
b
SPFI - glass surface
58 7705
44464 21028
c
SP correlation
PLL surface
d
SP correlation
glass surface
e
FCM buffer
FITC-H
EV size (nm), medium RI
f
FCM sample
g
NTA
Occurence
EV diameter (nm)
h
SP
FCM
NTA
Occurence (a.u., log10)

Fig S12: SPFI with CD63-GFP tagged EVs captured on two different surfaces. **a** Electrostatic capture on PLL coated glass coverslips (identical to Fig. 5**f**). **b** Capture on plasma activated bare glass. Small shifts in the distribution between **a** and **b** are visible, however, the two clusters of spots are clearly identifiable in either graph. **c** SP correlation plot on PLL. **d** SP correlation plot on glass. Once again, minor differences between **c** and **d** are visible, yet overall, the distribution is similar and the least square fit to the purple dots (solid line) used for stitching the SP contrasts had the same slope. **e** FCM measurement of EV sample buffer (PBS), see Fig. S14 for the FCM size calibration. **f** FCM measurement of the A431-GFP EVs (same data as Fig. 5**c** of the main text). **g** NTA measurement of the A431-GFP EVs. All data in this figure was taken with the same sample on the same day. **h** Comparison of the size distributions measured with SPFI, FCM and NTA. Histograms were generated from the data shown in **b**, **f** and **g**. For FCM, the buffer data was subtracted. All histograms were individually normalized to the bin centered at 130 nm to enable easy visual comparison. See text for a discussion on EV sizing with these three methods.

Fig. S13 shows additional data on CD63-GFP tagged A431 EVs which were purified using a 1 hr 10,000 g centrifugation step. Cells were cultured as described in the methods, a total of 135 ml of supernatant was spun at 400 g to remove cell debris and concentrated to 2 ml using 100 kDa molecular weight cutoff centrifugation filters (Amicon Ultra15 spun at 3,000 g for 30 min). The EV pellet was recovered, resuspended in 2 ml of PBS, spun again for 1 hr at 10,000 g and finally resuspended in 500 µl of PBS. No additional filtering was done.

The UC purified samples allowed us to further investigate the contrast switch of iSCAT on large EVs and identify the sizing limit in SP. Fig. 13**a** shows the SPFI plot after capture on PLL where we measured EVs with single pixel contrasts up to $\sqrt[3]{C_{combined}} \approx 0.7$. This was only marginally larger than the maximum $\sqrt[3]{C_{combined}} \approx 0.6$ found for SEC purified samples (c.f. Fig. S12**a**) and thus indicated that the limit for SP sizing with 740 nm was reached. Fig. 13**b** shows the correlation plot of both iSCAT contrast measurements, which clearly depicts the contrast switch in the 440 nm channel. Only very few particles that underwent the contrast switch at 740 nm (data not shown), and almost all of them showed irregular non-symmetric shapes suggesting there were plenty of aggregates. Fig. S13**d** shows a representative FOV of the post-incubation image measured in 440 nm, as well as exemplary spots measured in 440 nm (top row) and 740 nm (bottom row) whose coordinates can be matched with the arrowheads in Fig. S13**b**. The contrast switch in 440 nm suggests that EV sizes increase from the right to the left. The 740 nm measurement on the other hand shows roughly constant values (albeit at large variance) suggesting that it was close to the sizing limit. The spots with largest contrasts often showed irregular shapes (rightmost panel), likely indicating aggregation or collapse of EVs. The wealth of information present in the iSCAT image data suggests that EV sizing can be improved from the single pixel measurements performed in this manuscript by incorporating a full interferometric PSF model.

Fig. S13**c** shows the same EVs measured on a CytoFLEX-S FCM (see methods) for 60 s with 5 % of total events are plotted. Similar to the SEC purified sample shown in Fig. 5**c** of the main text, it was possible to detect a diffuse GFP signal on EVs above the detection threshold, however, the two populations found in SPFI could not be resolved. The FCM calibration was done using polystyrene beads in FCMPass software (see Fig. S14). We note that sizing of EVs between ca. 400 – 500 nm is restricted on the CytoFLEX-S due to Mie resonances (see flat region in the calibration curve of Fig. S14) which distorts the size distribution around this range after calibration and is thus not an accurate size measurement.

Fig. 13**e** shows NTA data, confirming large EVs > 300 nm in the sample. We note that this data was taken at reduced camera level 10 (see methods). Level 14, which was used for the SEC purified sample and allows to detect smaller vesicles, saturated the camera for EVs > 300 nm and led to a distortion of the size distribution. The slight shift in NTA peak compared to Fig. 5**d** is thus a measurement artifact due to the limited dynamic range of the NTA in single measurements which required us to lower the camera level and thus the sensitivity for small EVs.

The EVs were furthermore measured by negative stain TEM in Fig. S13**f** which confirmed the presence of both large vesicles around 500 nm as well as small ones below 50 nm diameters.

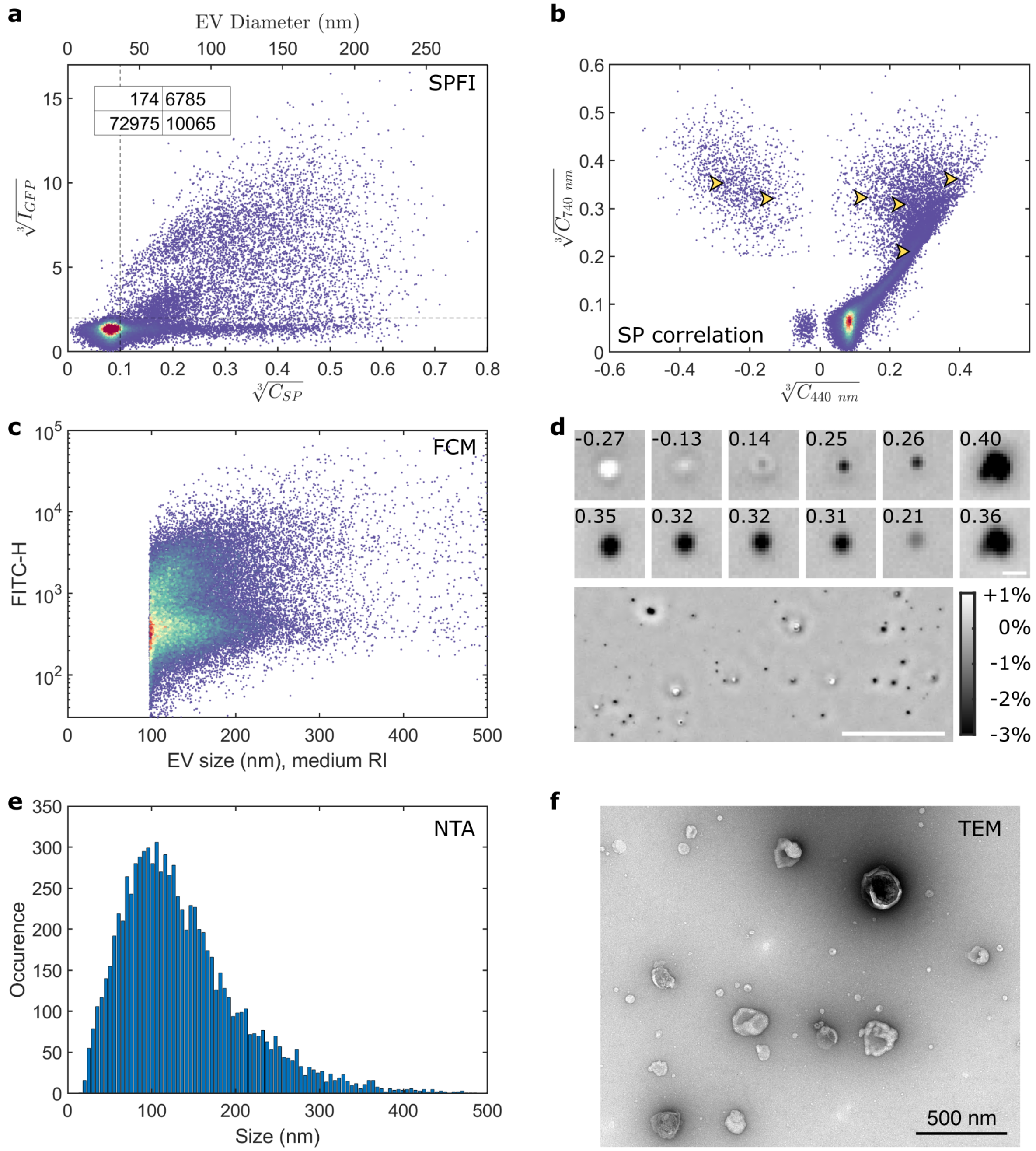

Fig S13: A431 CD63-GFP tagged EVs purified in a 10,000 g UC spin and measured in various techniques. **a** SPFI plot of the EVs captured on PLL coated glass. **b** iSCAT correlation plot of the two imaging wavelengths depicting the contrast switch in 400 nm. Arrowheads indicate the position of selected spots shown in **d**. **c** FCM data taken on a CytoFLEX-S, see Fig. S14 for the size calibration. **d** Exemplary post-incubation iSCAT data of the sample shown in **a** and **b**. The small panels correspond to the highlighted spots in **b** where the top row contains image data at 440 nm illumination and the bottom row the same spots imaged at 740 nm (scalebar 500 nm). The larger panel below shows a larger FOV imaged in 440 nm (scalebar 20 µm). The colorbar applies to all panels. **e** NTA data for the sample. **f** Negative stain TEM.

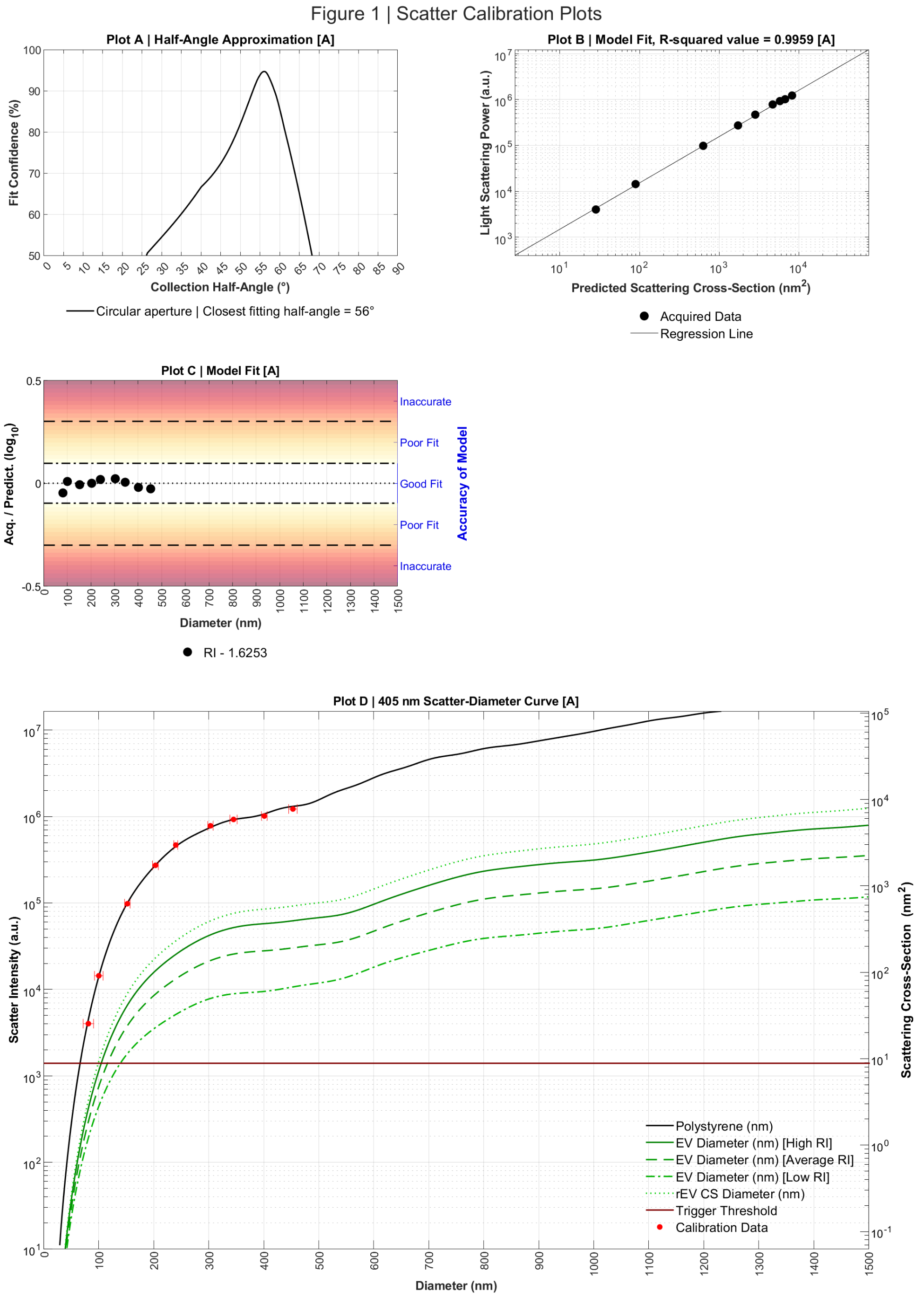

Fig S14: FCM calibration sheet exported from FCMpass v. 4.2.14.

### S10: Microscope objectives

All data presented in the main text were taken on a 100× oil immersion objective. However, we also tested a 60× objective to achieve higher throughput, Fig. S15. All datasets were collected with an additional 1.5× magnification lens (standard feature of the Nikon Ti2 that can be inserted into the light path with a manual rotary knob) as it was found that this produced higher contrasts for either objective lens. This resulted in full 1608×1608 pixel FOV sizes of (118 µm)$^2$ for the 100× objective and (197 µm)$^2$ for the 60× objective which thus yields a ~2.8 times greater imaging throughput. We found, however, that the raw iSCAT contrasts were reduced to about a quarter which corresponds to $\sqrt[3]{4} \approx 1.6$ times higher LOD with regards to particle diameter for 60× imaging in third root scaling.

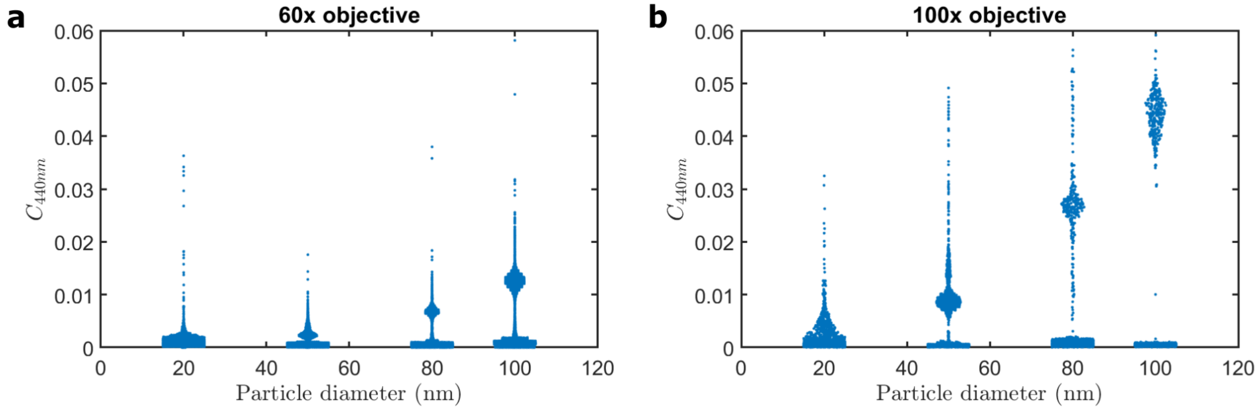

Fig S15: Comparison of iSCAT contrasts for silica nanobeads of known sizes. **a** Imaging with a 60× objective. **b** Imaging with a 100× objective. The image area with 60× objective is 2.8 larger, but at the cost of 4 times less contrast, and hence a ~1.6 times larger LOD with regards to particle diameter.
